# Human Peripheral Nerve-on-a-Chip on a Multiwell Microelectrode Array as a Scalable Preclinical Neurotoxicity Assay

**DOI:** 10.1101/2025.06.09.658615

**Authors:** Corey Rountree, Tyler Rodriguez, Ethan Byrne, Eva Schmidt, Erin Schwarzbach, Sabrina Dillon, Ally McCrimmon, Megan Terral, Monica Metea, Michael J. Moore, J Lowry Curley

**Affiliations:** 28bio Inc, New Orleans, LA; Preclinical Electrophysiology Consulting, LLC, Boston, MA; Tulane University, New Orleans, LA

## Abstract

Reliable human-relevant models of peripheral nerve function remain a critical unmet need in preclinical drug development, particularly for predicting neurotoxicity and bridging the gap to clinical translation. Here, we introduce a next generation Nerve-on-a-Chip, PNS-3D organoids, as a novel human-cell-based 3D peripheral nerve microphysiological system (MPS) that recapitulates key functional and structural features of native nerves, including long-distance axonal outgrowth, physiological myelination, and clinically translational population level electrophysiology. The platform integrates iPSC-derived human sensory neurons and primary human Schwann cells within a spatially organized 3D environment, coupled to a custom embedded electrode array that enables high-content, longitudinal, and clinically translatable functional assessments. As a proof of concept, we evaluated the platform’s predictive power using vincristine, a chemotherapeutic agent known to cause chemotherapy-induced peripheral neuropathy (CIPN). PNS-3D organoids captured dose- and time-dependent deficits in nerve conduction velocity, compound action potential amplitude, and axonal degeneration, with IC₅₀ values in line with human clinical exposures— outperforming traditional 2D cultures and *in vivo* benchmarks. Transcriptomic and morphological analyses further revealed neuron-specific degeneration consistent with axonopathy. These results validate the platform as a human-relevant, clinically translation, and scalable solution, enabling mechanistic safety assessment and drug discovery for neurotoxic and neuroprotective therapeutics.

## Introduction

Drug safety remains a primary clinical concern during the development of novel therapeutics, with neurological side effects commonly arising as off-target toxicities. Adverse clinical effects can include seizures, motor dysfunction, sensory deficits, or chronic pain, which severely impact patient well-being and frequently necessitate dose modification or discontinuation of treatment. Despite improvements in early screening for systemic toxicities, reliable pre-clinical platforms to detect or predict human-specific neurotoxicity remain elusive^1^. As a result, the ability to comprehensively evaluate the safety profile of therapeutic candidates, and the development of neuroprotective strategies, has been hampered by limitations in existing preclinical models.

Chemotherapy-induced peripheral neuropathy (CIPN) is a prime example of an unmet clinical need. CIPN is a common and often dose-limiting side effect of many frontline chemotherapeutics - including vinca alkaloids (e.g., vincristine), platinum-based drugs, and taxanes - with approximately 30-40% of treated patients developing peripheral neuropathy and neuropathic pain^2^. This condition not only erodes long-term quality of life in cancer survivors but can also necessitate treatment cessation in severe cases. Despite its prevalence and clinical significance, there are currently no approved preventive treatments for CIPN, although some are currently in clinical development^3,4^. Traditional rodent models have yielded important insights into CIPN mechanisms, but interspecies differences limit their predictive value for humans, and standard *in vitro* assays fail to capture the complex multicellular interactions and functional deficits that characterize human neuropathies^5,6^. These limitations have intensified the demand for advanced *in vitro* platforms to better predict neurotoxic side effects and to accelerate the discovery of neuroprotective therapies.

In this context, human microphysiological systems (MPS), or “organ-on-a-chip” models, have emerged as promising tools to bridge the translational divide. These *in vitro* systems combine tissue-specific architecture, dynamic environments, and primary or stem-cell-derived human cells to more closely mimic the structural and functional features of native human organs^7,8^. MPS have demonstrated value across organ systems, including the liver, heart, and kidney, and their utility for neurotoxicity screening is increasingly recognized. However, recreating the complex architecture and function of the nervous system *in vitro*, particularly the peripheral nervous system (PNS), remains a formidable challenge. Due to its complex three-dimensional (3D) architecture, unique electrophysiology, myelination dynamics, need for precise co-culture of neurons and glial cells, and limited availability of primary human cells, PNS-focused MPS have historically lagged other organ systems. Recent advances are closing this gap, as new platforms begin to recapitulate key features of human nerves *in vitro*, laying the groundwork for advanced models capable of clinical translational^9^.

Several models have employed microfluidics to recreate a model of the neuromuscular junction connecting human motor neurons to muscle fibers, demonstrating functional synaptic connectivity, providing a platform for studying neurodegenerative diseases such as amyotrophic lateral sclerosis in a dish^10^. Similarly, induced pluripotent stem cell (iPSC)-derived co-culture of motoneurons and Schwann cells have been used to replicate key aspects of nodes of Ranvier, thereby enabling investigations of hereditary neuropathies like Charcot–Marie–Tooth (CMT) disease in a complex biomimetic human environment^11^. Chung and colleagues advanced this space further by developing a fully 3D *in vitro* model of the PNS that integrates human Schwann cells and neurons within compartmentalized microfluidic chambers to guide axonal outgrowth and myelination under biomimetic conditions^12^. Their work demonstrated the feasibility of long-term coculture and functional evaluation in a closed system suitable for pharmacological testing.

Other groups have pioneered 3D engineered models of the PNS that illustrate the state of the art in this arena. Tissue engineering and biomaterials approaches have been leveraged to incorporate relevant cell types while enabling spatial patterning to recreate cell-cell interactions^13,14^. For example, both additive 3D manufacturing and scaffold-based electrospun fibers have been used to replicate peripheral nerve architecture from primary rodent cells and test functional recovery and myelin integrity following insult^15,16^. In one of the first examples of an all-human Nerve-on-a-Chip system, a combination of human iPSC sensory neurons and primary human Schwann cells produced millimeter-scale axonal outgrowth characterized by functional myelination^17^. Recreating the appropriate biology in dense 3D microfiber tracts also uniquely enabled the measurement of nerve conduction velocities *in vitro,* setting the stage for clinically translational electrophysiological endpoints. These efforts collectively highlight the growing consensus that, in order to achieve predictive translational outcomes effective PNS models must replicate both the structural and functional hallmarks of native nerves, including axon-glia interaction, 3D organization, and electrophysiological activity.

Previously, we published two case studies using our Nerve-on-a-Chip model to successfully track subtle pathological changes in nerve function following exposure to four common neurotoxic chemotherapeutics, though throughput was a limitation^18,19^. Here, we present the next generation model, PNS-3D organoids, designed to meet the need for a physiologically relevant, human-based system of peripheral neuropathy capable of longitudinal studies with significant increases in throughput. In addition, the improvements unlock higher-content data with additional clinically translational functional electrophysiological endpoints and multiplexing capabilities. The system incorporates iPSC-derived human sensory neurons and primary human Schwann cells within a spatially organized 3D microengineered environment integrated within a custom embedded electrode array (EEA) plate, yielding an all-human nerve tissue construct that structurally and functionally resembles a native peripheral nerve. This biomimetic model recapitulates the formation of myelinated axon bundles and supports electrophysiological recordings analogous to clinical nerve conduction studies. The model is also compatible with multiplexed imaging and cell viability assays, as well as transcriptomic profiling, allowing comprehensive assessment of neuronal health and pathology under different treatment conditions. We demonstrate the utility of PNS-3D organoids by testing vincristine – a neurotoxic chemotherapeutic well known for causing CIPN – and observing dose and time-dependent electrophysiological impairments and cellular changes that mirror *in vivo* and clinical findings. Taken together, these capabilities position our platform as a valuable drug screening tool for CIPN, offering a new means to evaluate the neurotoxicity of therapeutic drug candidates and potential neuroprotective therapies in a human-specific context.

Beyond CIPN, the versatility of the model also lends itself to broader applications, including mechanistic studies of hereditary demyelinating neuropathies (such as CMT) and investigations of chronic pain pathophysiology, where improved *in vitro* human models of the peripheral nerve are likewise critically needed.

## Results

### Construct Biofabrication and Assay Endpoints

Initially, human iPSC derived sensory neurons (hNs) and primary human Schwann cells (hSCs) are cultured in 2D flasks to encourage proliferation and remove non-viable cells following thaw from cryopreservation (Figure 1A). After 24 and 48 hours for hNs and hSCs respectively, cells were passaged and co-cultured for four days to encourage organoid formation. Finally, organoids were transferred to circular growth regions of individual constructs within a specially designed 24-well plate. These are grown in standard cell culture conditions demonstrating continuous neurite outgrowth across time with neurites observed down the entirety of the channel after 21 days. Based on additional criteria to be discussed throughout this manuscript, day in vitro (DIV) 42, measured post-spheroid incorporation, is used as the timeline to constitute mature samples ready for testing.

**Figure 1.**
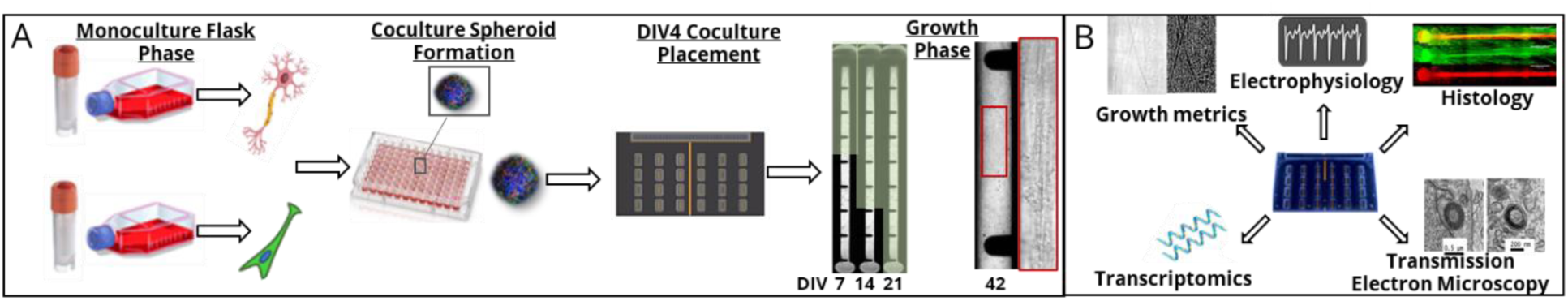
Biological fabrication of PNS-3D organoids. A) Cell culture and biomanufacturing workflow from left to right in which hIPSC-derived sensory neurons and primary human Schwann cells are thawed in monoculture, for 24 and 48 hours, respectively. The cells are lifted to form coculture organoids in a U-bottom plate for 4 days. Each organoid is then placed in individual constructs for a 42-day growth period. B) Endpoints within the EEA plate can be multiplexed allowing for functional and morphological metrics including electrophysiology, growth, immunohistochemistry, transmission electron microscopy and transcriptomics.

### Design and Manufacturing of a Custom Embedded Electrode Array Platform for *In Vitro* Electrophysiological Recordings

To enable physiologically relevant electrophysiological measurements across long distances within an *in vitro* peripheral nerve model, we developed a custom microelectrode array (MEA) platform specifically tailored for peripheral nerve conduction studies. Existing commercial MEA systems prioritize high-density electrode placement optimized for local field potential mapping and network bursts but are poorly suited for detecting compound action potential (CAP) propagation over centimeter-scale distances. These constraints necessitated the design of a new system that could support spatially resolved stimulation and recording over extended axonal tracts, thereby allowing velocity, excitability, and spatially dependent effects to be assessed with high precision.

The MEA was fabricated using a flexible polyimide-based printed circuit board (PCB) substrate, enabling high-resolution electrode patterning while maintaining compatibility with standard laboratory equipment (Figure 2A). Each of the 24 wells incorporates 10 soft gold-coated electrodes (50 μm diameter) and two large reference electrodes (1.2 mm diameter) configured for low-noise differential recordings. Electrode traces (100 μm wide) connect to a 300-pin Samtec connector on the plate’s edge, supporting repeated, noise-resistant interfacing with an electrophysiology system. The overall footprint conforms to ANSI/SBS microplate standards, and the translucent substrate permits high-content inverted microscopy.

**Figure 2.**
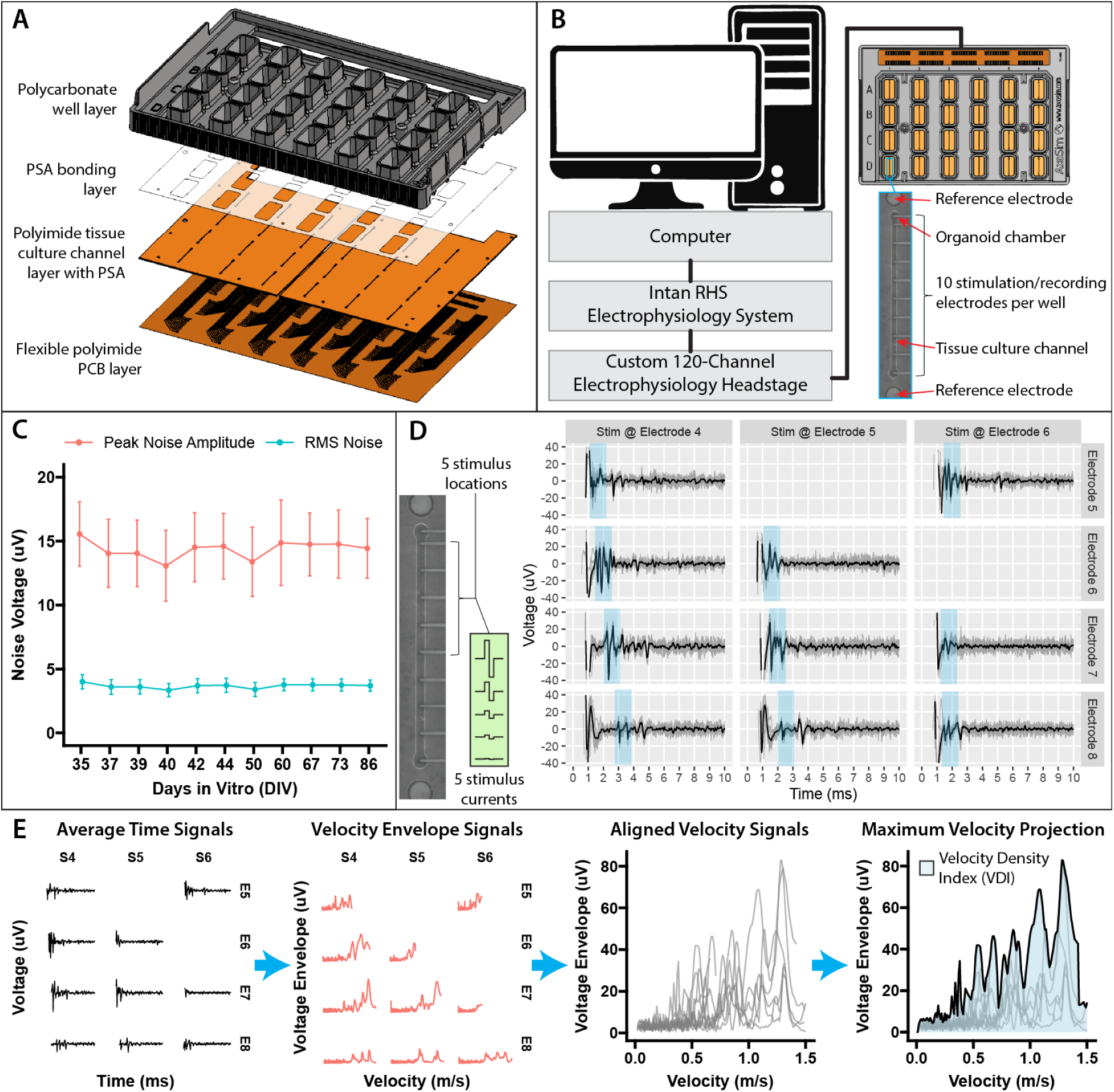
Design, Integration, and functional characterization of the Embedded Electrode Array (EEA) plate for multichannel neural recording and analysis. A) Computer aided design rendering of the multilayer EEA plate design (from bottom to top): flexible polyimide-based printed circuit board (PCB), polyimide-based tissue culture channel layer with pressure-sensitive adhesive (PSA) for lamination, additional PSA layer, and final injection-molded polycarbonate structure for well reservoirs and structural integrity. B) Electrophysiology setup schematic with EEA plate interfacing with a custom 120-channel headstage enabling simultaneous recording from 12 wells. Headstage connects to an Intan RHS recording system. Inset (bottom right) highlights the microchannel design (200 μm width, 500 μm depth) guiding neuronal growth across 10 mm, with ten 50 μm-diameter stimulation/recording electrodes spaced at 1 mm intervals and flanking reference electrodes. C) Stability and sensitivity demonstrated through noise performance data from N = 18 plates, showing consistent low root-mean-square (RMS) noise (blue) and peak noise amplitude (red) across time. D) Electrophysiological recordings and stimulation paradigm. Left panel shows stimulation across five electrodes at varying current amplitudes (1–48 μA). Right panels display neural responses recorded at four electrodes (rows) in response to stimulation at three different sites (columns), with each trace representing the mean (black) of 12 stimulation trials (grey). Early response components highlighted (blue) to indicate conduction timing across the array. E) Data transformation pipeline from raw recordings to the velocity density index (VDI). Raw time-domain signals rectified and transformed into velocity space (second panel), aligned by velocity (third panel), and aggregated with maximum across traces at each velocity (black trace, fourth panel). VDI is calculated as the area-under-the-curve of this maximum.

A laser-cut, 500 μm-thick polyimide culture layer is affixed atop the PCB to spatially constrain axonal growth. Each well contains a central microchannel (200 μm wide) flanked by 700 μm organoid chambers to house organoids and direct axonal projection across the embedded electrodes. This layer is bonded to the PCB using a pressure-sensitive adhesive (PSA, 25 μm thick), itself patterned to match the microchannel geometry. A second PSA layer defines the fluidic well footprint, and a final injection-molded polycarbonate shell forms the uppermost layer, providing structural rigidity and individual reservoirs for media exchange. The polycarbonate layer has a custom lid design that allows electrophysiological recordings without lid removal, thereby maintaining sterility throughout experimentation.

### System Integration, Signal Acquisition, and System Performance

Signal acquisition and stimulation were implemented using the Intan RHS platform— originally designed for *in vivo* recordings in freely moving animals (Figure 2B). We engineered a custom 120-channel headstage using Intan’s integrated circuit (RHS2116) amplifiers and analog-to-digital converters, allowing full coverage of half the plate (12 wells) with switching between top and bottom halves. Communication with the Intan RHS controller occurs via USB and is orchestrated through a custom-developed MATLAB interface. This system automates the stimulation protocol by parsing Excel-defined stimulation parameters (location, polarity, current amplitude, pulse width, frequency, and number of trials), enabling high-throughput, reproducible experiments. The software archives stimulation data by plate and recording session, and supports paradigms including current ramps, spatial analyses, and activity-dependent slowing protocols—each aimed at extracting clinically and biologically relevant metrics.

System performance was benchmarked over long-term culture (day 35–86) across 18 independent plates (Figure 2C). Noise characterization demonstrated stable low-noise conditions compatible with both single-spike detection and compound action potential analysis. RMS and peak amplitude measurements were consistent across timepoints and wells, confirming suitability for longitudinal studies and cross-comparative experimental designs.

### Functional Testing and Endpoint Analysis

After periods of cell culture from DIV 35-86 we validated the system’s ability to detect stimulus-evoked CAPs (Figure 2D). Signals were reliably distinguished from background using threshold-based detection (6σ above mean noise) and further refined with machine learning approaches. The standard assay involved stimulation at five discrete positions (0.45–4.45 mm from the organoid), enabling analysis of both proximal and distal activation. Current ramps from 1 to 48 μA were applied to identify activation thresholds.

This platform supports extraction of both classical and novel electrophysiological endpoints. Clinically relevant parameters include Nerve Conduction Velocity (NCV) calculated using known distances between stimulation and recording sites and response latencies and CAP Amplitude determined via peak-to-peak (max–min) signal amplitude, to be discussed later.

In addition, we developed novel *in vitro* metrics tailored to the system (Figure 2E): Number of Peaks reflects different subpopulations of neurons firing at different velocities, quantified as the number of deflections above 6σ. Maximum Velocity Projection (MVP) is a composite waveform derived by converting time-domain traces into velocity space and taking the envelope maximum across all channels, and Velocity Density Index (VDI) represents the area under the MVP curve.

Taken together, this scalable, multi-layered MEA platform enables long-distance high-content *in vitro* electrophysiology across biomimetic human nerve tissues

### Co-culture Organoid Dynamics across Cellular Densities

First, the ability of neuronal cells and Schwann cells to form co-culture organoids was assessed based on the percentage of plated organoids which successfully formed (Figure 3A). At the end of the organoid formation period (DIV4), organoids were imaged in phase at 10X magnification. Organoid area (µm^2^) was measured for each co-culture density. 100% of organoids across all conditions successfully formed. Organoids must remain <1000 µm in diameter to fit into the circular chamber of the construct, and all of the conditions satisfied this criterion, with all organoid remaining below 1000 µm. Organoid size was assessed across varying organoid densities to better understand the interactions between neurons and Schwann cells, and the resulting effect of increasing the number of each cell type on organoid size. Average diameters were measured to be 536 µm, 614 µm, 788 µm and 858 µm for neuronal to Schwann cell ratios 20,000:10,000, 30,000:15,000, 40,000:20,000, and 50,000:25,000, respectively. To assess the advantages of different organoid densities, a subset of conditions were cultured in a 3D hydrogel for 4 days and imaged to evaluate neurite outgrowth (Supplemental Figure 1). Qualitatively, outgrowth remained relatively consistent across neuronal densities in monoculture. However, the length of neurites appeared to increase with increasing densities of Schwann cells. A single density of 50,000 hSNs and 25,000 hSCs per organoid was chosen for subsequent experiments to maximize the number and density of nerve fibers for electrophysiological recordings.

**Figure 3.**
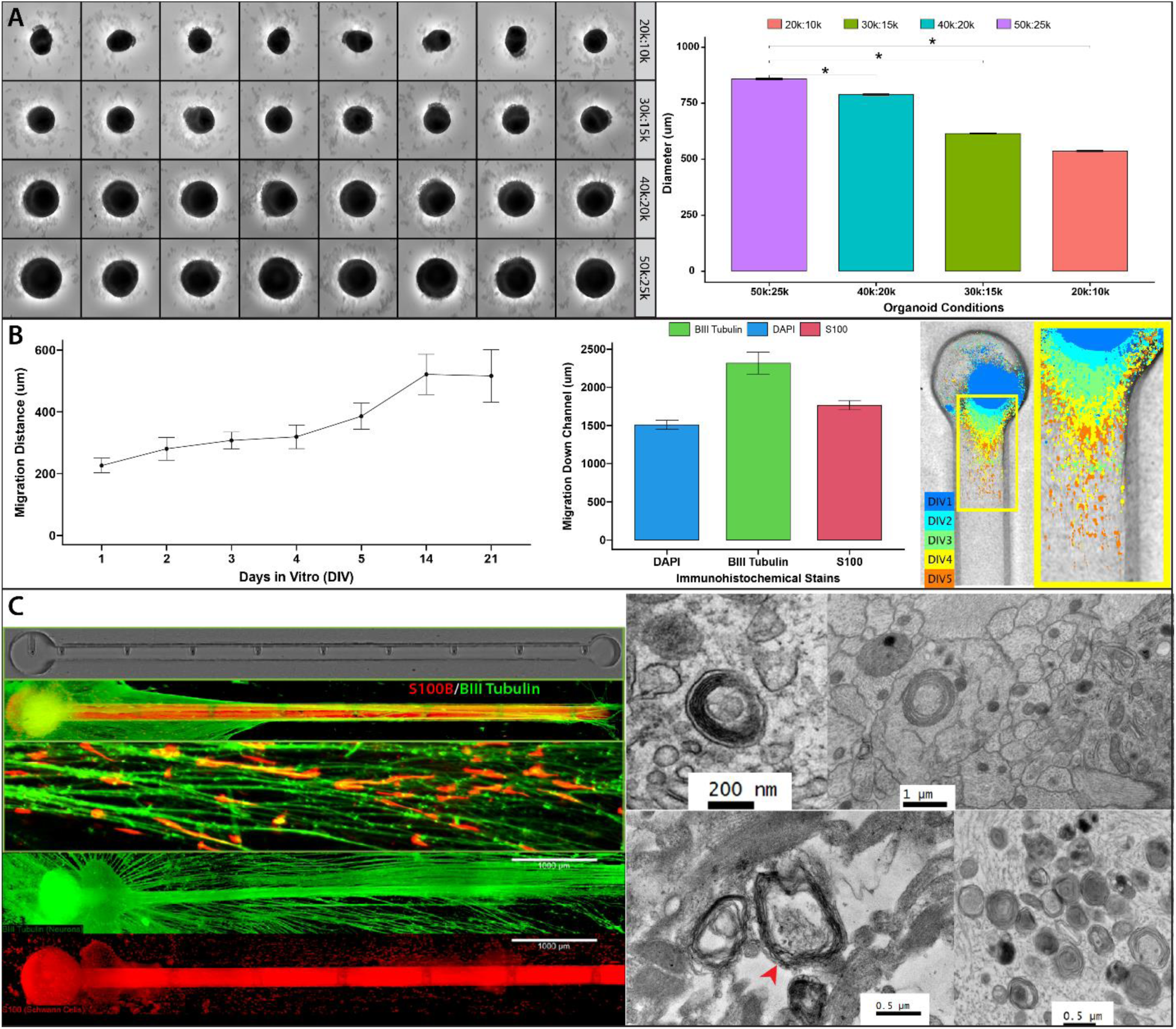
Biological characterization of PNS-3D organoid co-culture model. A) Density optimizations of coculture organoids formed from iPSC derived sensory neurons and human primary Schwann cells to model the peripheral nerve. Transmitted light images of DIV 4 organoids were taking to measure areas of various ratios of hSNs to hSCs from top to bottom: 20k:10k, 30k:15k, 40k:20k, and 50k:25k and the corresponding quantified diameters of each ratio tested (right). B) Schwann cell migration was tracked for five consecutive days, two weeks, and three weeks using fluorescent microscopy of Qtracker-labeled Schwann cells (left). Because Qtracker can wash out of proliferating cells, DIV7 samples were fixed and stained to track cell migration with Beta III Tubulin for neuronal outgrowth S100 for Schwann cells, and DAPI (middle). Visualization of Qtracker-labeled Schwann cells for DIV 1-5 (right). C) To further qualitatively confirm outgrowth and Schwann cell migration DIV 49 samples were stained and imaged. Single stains and overlayed images allow visualization of Schwann cell alignment with neurons, ideal for electrophysiology and a biomimetic model. D) Myelination was present in DIV 49 samples as lamellar bodies containing an axon (red arrow).

### 3D Neurite Outgrowth, Schwann Cell Migration, and Myelination

When cultured inside of the full system, neurite outgrowth reached the end of the constructs after 21 days and Schwann cell migration was quantified along the same period (Figure 3B). Live Schwann cells were incubated with a Qtracker labeling kit in monoculture before organoid integration to allow non-invasive, longitudinal Schwann cell migration tracking via fluorescent microscopy. Schwann cells were observed to migrate throughout the culture period with significant initial proliferation observed, reaching approximately 511 µm and 500 µm at days 14 and 21. Because rapid proliferation of Schwann cells can dilute the Qtracker labels, a subset of samples were fixed and stained via immunohistochemistry (IHC) at DIV7 to provide a more accurate migration estimation. The DIV7 IHC results showed that neurite outgrowth preceded Schwann cell migration, with neuronal outgrowth reaching 2,319 µm down the channel while Schwann cells migrated 1,562 µm. After the full 49 days in culture, both were observed down the entire 10 mm length of the channel (Figure 3C). Neurite outgrowth density also increased with time, filling the entirety of the channel, though overgrowth was observed outside of the channel. Schwann cells were seen to have migrated and continued proliferating throughout culture within the 3D channel with some moving over the top of the channel. The majority of Schwann cells observed were associated closely with neurites, with most directly aligned with the neurites though some were still circular in morphology.

To visualize biological ultrastructures, fixed cross-sectional images were obtained with transmission electron microscopy, demonstrating randomly distributed Schwann cells throughout the cross sections. Many, though not all, were associated with axons, as also observed through IHC. The majority of myelin was present as lamellar bodies containing more solid structures. This sometimes appeared in a disorganized fashion but often as loosely packed, concentric rings with small central spaces. Some myelin was present in faux sheaths of concentric circles of variably packed lamellae arranged around an axon with examples also visualized having an empty central area.

### Highly Parallel Electrophysiological Stimulation and Recording

Using a custom embedded electrode array (EEA) electrophysiology system, up to 12 cultures can be tested in parallel with current-based electrical stimulation. The system was designed to be scalable and automated with the throughput of approximately 10,000 individual electrode recordings (each recording containing 12 trials of stimulation at 1 Hz) per hour. A typical recording session involves stimulation at multiple sites with a stimulation current ramp (1 to 48 µA) while recording the CAPs at the organoid body as well as along the axons to fully characterize the sensitivity of the neuronal cultures to electrical stimulation. As described previously, where reported, electrophysiological recordings are shown as raw signals (µV) or processed into VDI (% normalized to the vehicle control) to quantify the population-level response to electrical stimulation. Voltage can be measured and compared based on the distance separating the recording electrode from the organoid, allowing quantification of changes to the whole nerve, and distinct changes near neuronal cell bodies or closer to axon endings (Figure 4 A & B). Results across time demonstrate consistency in electrophysiology, with an increase in higher voltage responses across time. A stimulation current ramp (1-48 µA) was tested at each site before and after exposure to a reversible blocker, 150 mM potassium (150 mM) to shut down the physiological gradient of sodium-potassium pumps, thereby stopping action potential generation and overall neural signaling. Results show the cessation of responses compared to baseline prior to exposure (Figure 4C). After washout, responses returned, demonstrating the biological nature of the functional recordings. In a separate experiment, VDI was calculated for whole nerve, proximal, and distal similar to Figure 4A. VDI values started at 7.94 µV*ms (sem = 2.73) for whole nerve, 6.47 µV*ms (sem = 2.2) for proximal nerve, and 4.17 µV*ms (sem = 2.04) for distal nerve, respectively, and remained consistent across time for all spatial locations (Figure 4D). Similarly, at dosing day 7 in the same experiment, VDI values were shown to increase with increasing stimulus current as expected for a multi-fiber neural tract rising from 0.1 µV*ms (sem = 0.275) at 1 µA to 8.85 µV*ms (sem = 0.727) at 24 µA before reaching a maximum of 14 µV*ms (sem = 2.93) at 48 µA (Figure 4E). As VDI values demonstrated a plateau after 48 µA of stimulation, future results were generated with a maximum stimulation current of 48 µA. The assay can be performed at multiple timepoints before, during, and after drug exposure to fully characterize the scale and timing of pharmacological effects.

**Figure 4.**
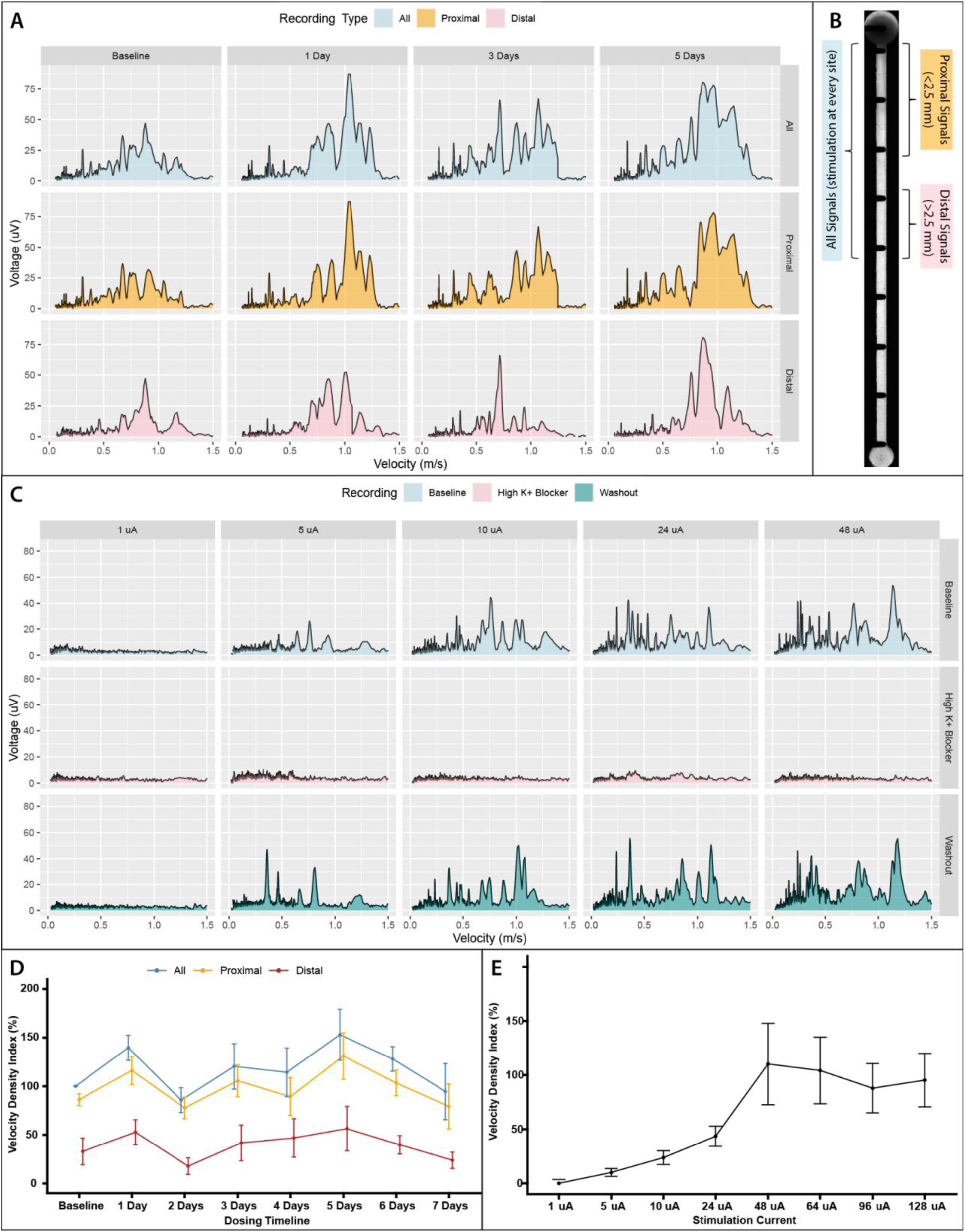
Electrophysiological data analysis examples and demonstration of biological responses. **(A)** Maximum velocity projection (MVP) profiles for a representative vehicle-treated sample over five days. Each column represents a different timepoint, and each row displays responses derived from different stimulation conditions: all stimulation sites (top), proximal stimulation only (middle; stimulation in first 2.5 mm of channel), and distal stimulation only (stimulation past 2.5 mm of channel). MVP traces (voltage vs. velocity) demonstrate progressive strengthening of responses across all components over time. **(B)** Schematic of the stimulation layout within a single EEA channel, illustrating how MVPs are generated from different subsets of stimulation sites: *all signals* include responses to all five sites, *proximal* includes stimulation at electrodes <2.5 mm from the organoid, and *distal* includes stimulation at electrodes >2.5 mm from the organoid. **(C)** Functional validation of stimulus-response sensitivity and pharmacological modulation. Columns represent increasing current amplitudes (1–48 μA); rows represent different recording conditions. Baseline recordings (top row) show graded increases in response magnitude with increasing current, with activation thresholds typically observed at 5 μA. Application of a reversible conduction blocker (150 mM K⁺; middle row) results in complete loss of neural responses across all current levels, confirming signal origin from active neuronal conduction. Recordings taken 30 minutes post-washout (bottom row) show restoration of neural responses, validating reversibility. **(D)** Quantification of the velocity density index (VDI) across one week in vehicle-treated samples, separated into *all*, *proximal*, and *distal* stimulation conditions. While some day-to-day variability is observed, VDI values remain elevated and stable, indicating sustained functional responses. **(E)** Quantification of VDI responses from (C), demonstrating a strong dependence on stimulation current. VDI increases steeply between 5–48 μA and plateaus at higher current levels (48–128 μA), suggesting saturation of responsive neuronal populations.

### Clinically-Relevant Metrics of Toxic Neuropathy Following Vincristine Exposure

As a proof of principle, we used vincristine-induced peripheral neuropathy to demonstrate the model’s effectiveness as a neurotoxic screening platform. Human iPSC cultures were grown for 42 days before dosing with various concentrations of vincristine (VN), ranging from 0.1 nM to 1000 nM. We performed daily electrophysiological recordings on the same samples over a 7-day dosing regimen. Each construct was electrically stimulated at multiple electrode positions along its length, enabling comprehensive mapping of population-level electrophysiological responses. These analyses revealed a clear dose- and time-dependent decline in nerve function, particularly pronounced at higher vincristine concentrations where functional responses rapidly declined within the first few days (Figure 5A). To quantitatively track these progressive changes, we computed IC_50_ estimates at each time point (N = 6 samples), revealing increasingly potent effects over time and confirming the cumulative neurotoxicity of vincristine (Figure 5B). Results indicated significant decreases in activity at concentrations of 10 nM vincristine and higher. Even after 7 days, 0.1 nM VN did not show a significant decline while effects for 1000 nM were seen as early as 16 hours of dosing. The IC50 values remained at or above 23.9 nM for the first 48 hours. A decrease in IC50 from 6.68 nM to 1.23 nM was seen from dosing DIV 3 to DIV 7, respectively.

**Figure 5.**
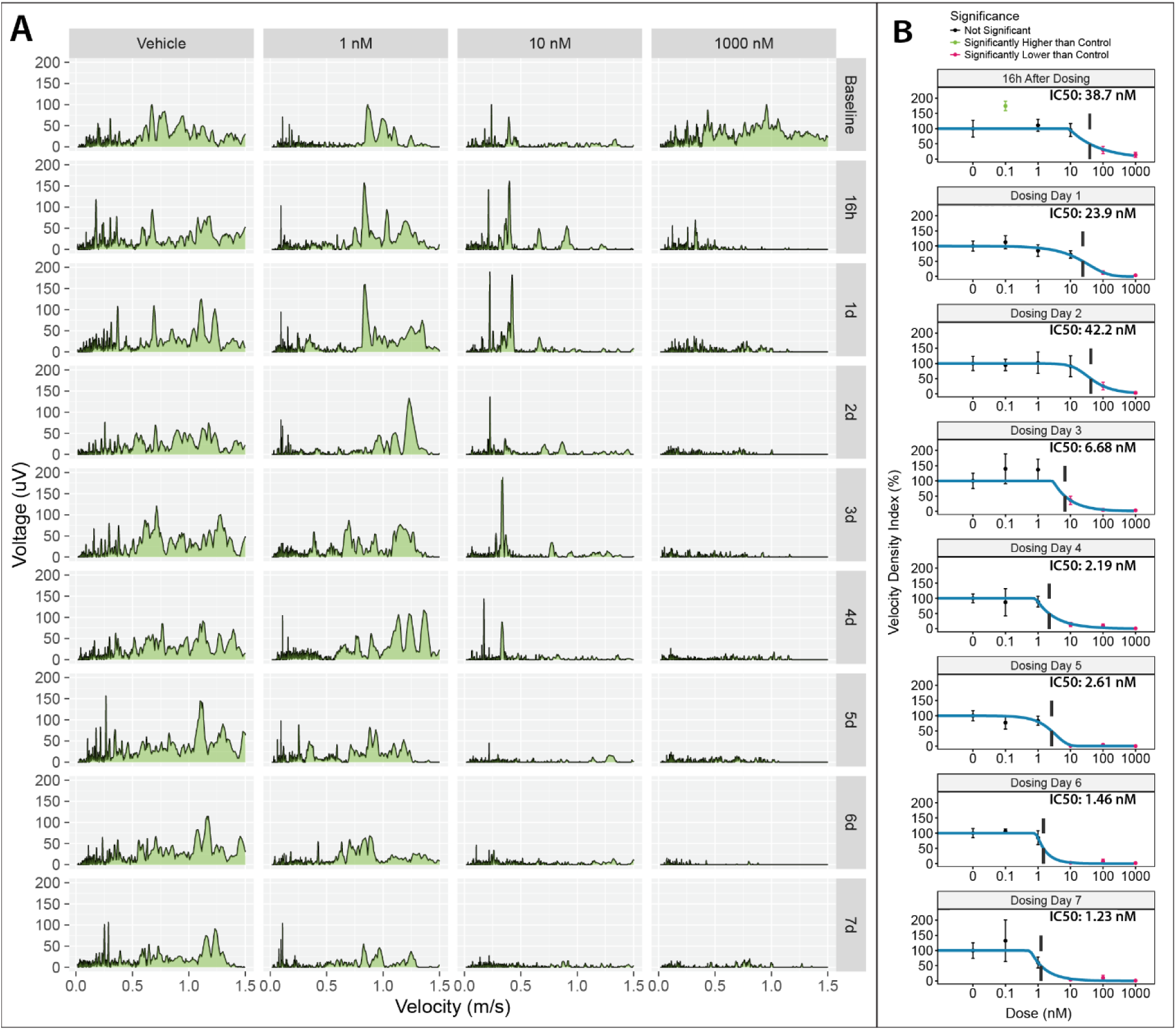
Prolonged application of vincristine induces dose-dependent toxic neuropathy. **(A)** Representative maximum velocity projection (MVP) profiles from individual samples exposed to increasing concentrations of vincristine over time. Columns correspond to different vincristine concentrations (0, 1, 10, and 1000 nM), and rows represent successive timepoints spanning baseline through 7 days of treatment. Each panel plots voltage versus conduction velocity, with the velocity density index (VDI; area under the curve) highlighted in green. Vehicle-treated samples (left column) maintain stable and robust activity across all timepoints. Samples treated with 1 nM vincristine retain strong signal responses, though subtle slowing emerges by days 5–7. At 10 nM, activity is initially strong but exhibits clear slowing and amplitude reduction by day 5. The highest dose (1000 nM) results in near-complete signal loss within 16–24 hours, despite robust baseline activity. **(B)** Quantified population data corresponding to (A), showing longitudinal VDI measurements across all wells for each dosage condition. Each panel displays a dose–response curve with an IC₅₀ fit (blue line), the calculated IC₅₀ value (dotted black line), and significance markers relative to vehicle controls (green: increase, red: decrease; *p* < 0.05). IC₅₀ values start relatively high (∼40 nM) in the early treatment period but progressively decrease over time, indicating increasing sensitivity to vincristine neurotoxicity. All data represent longitudinal measurements from the same samples over time, allowing within-sample tracking of neuropathy. Error bars represent standard error of the mean (SEM).

A separate cohort of samples were dosed at lower concentration ranges (0.01 nM to 100 nM) for 7 days to assess the repeatability of the 7-day IC50 calculation (Figure 6A). The 7-day VDI measurement were found to be closely aligned (IC50 = 1.03 vs 1.23 nM) with the measurements in Figure 5B. Further data analysis allowed for the measurement of additional clinically relevant electrophysiological metrics from compound action potential (CAP) recordings. These metrics included CAP amplitude, NCV, and number of responses (Figure 6A), each having distinct IC50 values following 7 days of dosing with VN. In addition, the MVP signals were broken down into velocity bins, partitioning the VDI into slow (0–0.5 m/s), medium (0.5–1.0 m/s), and fast (1.0–1.5 m/s) bins to highlight shifts in conduction profiles (Figure 6B). Across velocity bins, IC50 values demonstrating unique dose and time dependent changes in each population. High doses of vincristine caused progressive loss of fast and medium velocity responses after three days dropping from 15.7 and 10.1 nM at day 1 to 1.5 and 1.55 nM (Figure 6C) at day 3. However, IC50 values of slow velocity responses saw a slower decline from 32.3 to 8.34 at days 1 and 3. As described in Figure 4A-B, the VDI can also be spatially binned into proximal (<2.5 mm) and distal (>2.5 mm) groups, allowing comparisons of spatial function (Figure 6D). IC50 values for VDI values recorded for distal locations started lower at 3.02 nM compared to 22 nM for proximal. Subsequently, proximal IC50s demonstrated a decrease to 3.02 nM after 5 days, consistent with axonopathy.

**Figure 6.**
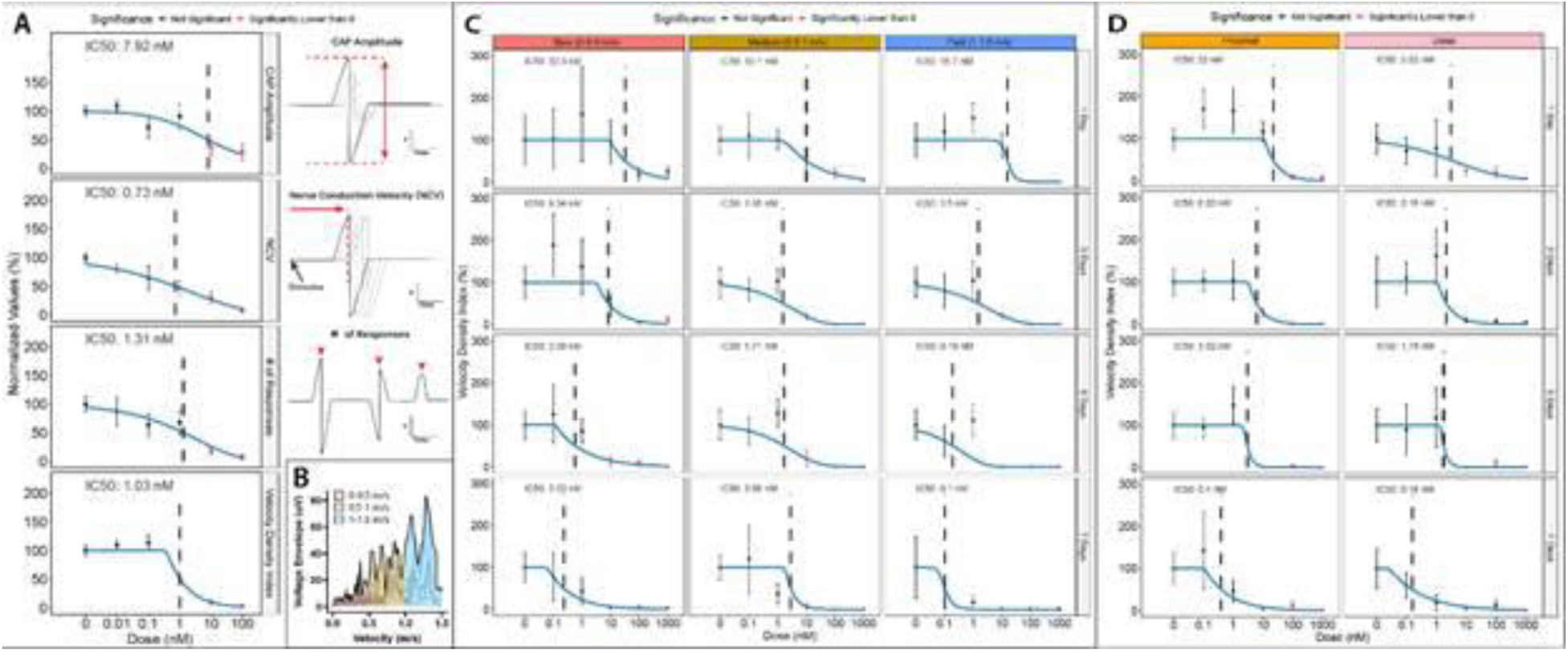
Additional clinically translation electrophysiological endpoints enable mechanistic insights. (A) This panel presents additional electrophysiological metrics from a follow-up experiment assessing vincristine-induced neurotoxicity at Day 7 of dosing, using a refined dose range (0.01–100 nM). The top plot shows the compound action potential (CAP) amplitude, measured peak-to-peak, with the IC₅₀ curve fit overlaid in blue, the calculated IC₅₀ value indicated with a black dotted line, and statistically significant reductions from vehicle controls marked in red. The second plot displays the nerve conduction velocity (NCV) of identified responses. The third plot presents the number of detected CAP responses, reflecting how vincristine dosage impacts different populations of sensory neurons. The bottom panel shows the velocity density index (VDI) data for this experiment, which closely aligns with findings from the prior study (see Figure 5B, bottom panel), reinforcing the VDI as a sensitive and reproducible readout of electrophysiological function. Graphical representations are included to the right of the first three metrics to explain the specific measurements. (B) A schematic representation illustrates the process of velocity binning used to decompose the MVP signal into distinct conduction velocity ranges. The total VDI is segmented into slow (0–0.5 m/s), medium (0.5–1.0 m/s), and fast (1.0–1.5 m/s) bins, with the VDI (area-under-the-curve) calculated separately for each region to capture bin-specific changes in neural conduction. (C) This panel quantifies the velocity-binned VDI across four timepoints (Days 1, 3, 5, and 7) shown in rows, with each column representing a different conduction velocity bin. Each panel includes an IC₅₀ curve and the associated value. Over time, the fast-conducting bin shows a progressive and near-complete loss of signal by Day 7 at all but the vehicle condition, while medium and slow bins exhibit more gradual reductions, primarily at higher vincristine concentrations. (D) Shown here is the quantification of spatial binning, as previously introduced in Figures 4A and 4B. Columns represent different spatial segments along the EEA channel, from proximal to distal, while rows indicate dosing timepoints. The IC₅₀ values for each spatial bin reveal a consistent trend of greater sensitivity in the distal regions, suggesting that vincristine induces a length-dependent degeneration pattern consistent with axonopathy, rather than a global neuropathy.

The currently configured PNS-3D organoids can also be multiplexed to provide a more complete profile of neurotoxic effects on peripheral nerve cultures. Each culture can be examined via brightfield microscopy to correlate spatial morphology with the functional, spatially localized electrophysiological data (Figure 7). Similar to functional changes, exposure to VN led to dose and time dependent changes in the nerve morphology (Figure 7 A-B) that can be captured using automated image analysis to quantify the nerve fibers and fragmentation of cells as separate binary masks. The data in these binary masks can be quantified to extract morphology metrics including fiber count, fiber length, and degeneration index (Figure 7 C-E). Degeneration index represents the fragmentation of cells due to toxicity, measured as the number of small particles in the fragment masks (see Figure 7 A-B). Overall, the nerve morphology metrics display a slower progression of neuropathy compared to the functional losses, with significant but not severe toxicity at day 2 contrasted to the complete loss of function for the equivalent time points (Figure 5B). After 7 days of dosing however, the IC50s for all growth metrics aligned well with the electrophysiological IC50s.

**Figure 7.**
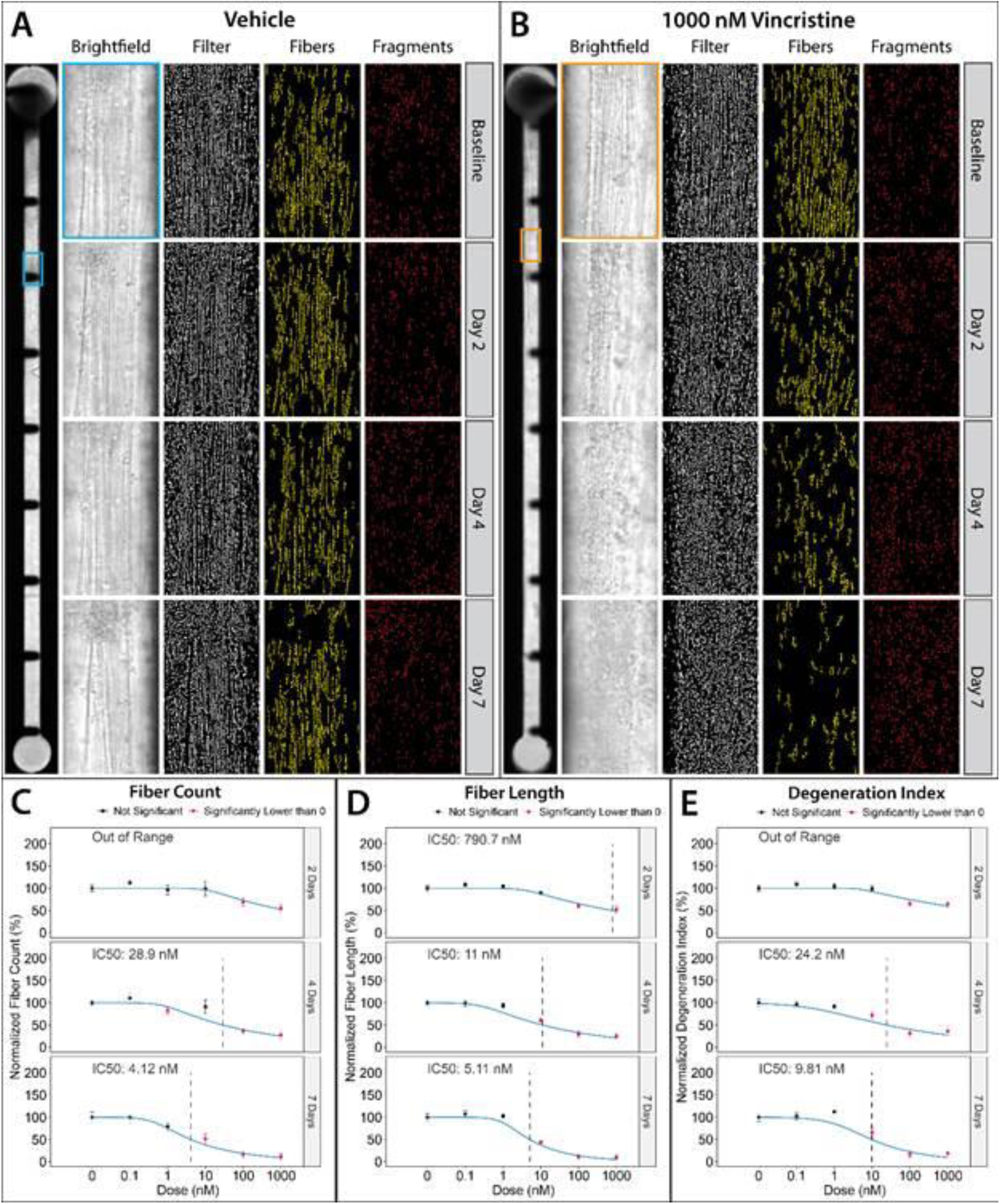
Axonal degeneration after VN exposure demonstrated unique dose and time dependent toxicity. (A) Representative brightfield imaging and morphological analysis of a vehicle-treated sample over a 7-day period. The leftmost panel displays a full-channel view of the embedded electrode array (EEA) at baseline, showing the organoid positioned at the top of the channel and the array of 10 stimulation/recording electrodes spaced along its length. A blue box indicates the region of interest, which is expanded in the adjacent panels. The second column presents high-resolution brightfield images across four timepoints (baseline, 2 days, 4 days, and 7 days) capturing neurite outgrowth and structural changes. The third column shows the output of a custom image filter that generates binary masks of brightfield images to highlight neurite structures. The fourth column displays the fiber mask, a refined output that filters for fibrous objects based on an aspect ratio ≥ 0.9 and a minimum length of 25 pixels. The fifth column presents the fragment mask, isolating smaller non-fibrous elements defined as objects with an area ≤ 25 pixels. Throughout the 7-day period, vehicle-treated cultures exhibit relatively stable morphology across all metrics, with consistent fiber and fragment representation. (B) Parallel image series from a culture treated with 1000 nM vincristine. In contrast to the vehicle condition, vincristine exposure results in a progressive loss of visible neurite fibers in the brightfield images and a concurrent increase in small cellular fragments. This trend is substantiated by the fiber masks, which reveal decreasing fiber count and length over time, and by the fragment masks, which show a marked increase in the number of small, fragmented structures. (C–E) Quantification of structural degeneration metrics was performed across *N* = 10 samples per condition. Fiber count (C), calculated from the number of objects in the fiber mask, exhibited a sharp decline at higher vincristine concentrations by Day 2, with IC₅₀ values clearly defined at Days 4 and 7. Average fiber length (D) also decreased in a dose- and time-dependent manner, with IC₅₀ values appearing as early as Day 2 and continuing to drop with prolonged exposure. The degeneration index (E), derived from the proportion of small objects identified in the fragment mask, mirrored these trends showing significant increases in fragmentation at early timepoints and IC₅₀ values measurable by Days 4 and 7. Together, these metrics provide a complementary and quantitative view of vincristine-induced neurite damage, capturing both loss of structural integrity and emergence of cellular debris over time.

To better understand the underlying molecular mechanisms, we also conducted bulk RNA sequencing on PNS-3D nerves collected at the end of the 7-day dosing period from vehicle-treated samples and two vincristine concentrations: 1 nM (around the IC50 threshold) and 10 nM (above IC50). Differential expression analysis (DESEQ2) revealed significant downregulation of genes involved in sensory neuron identity and electrophysiological function (42 of 157), alongside a marked upregulation of pro-apoptotic genes (10 of 19), particularly at 10 nM (Figure 8)^20–22^. In contrast, Schwann cell-associated gene expression remained largely unchanged (3 of 40), with markers like S100B showing no significant difference (Figure 8D)^21,23–26^. Notably, key sensory neuron markers such as neurofilament heavy chain (NEFH), PIEZO2, Nav1.6 (SCN8A), and Nav1.7 (SCN9A) were significantly reduced, while stress-associated channels like TRPV1 and Nav1.9 (SCN11A) were upregulated^27,28^.

**Figure 8.**
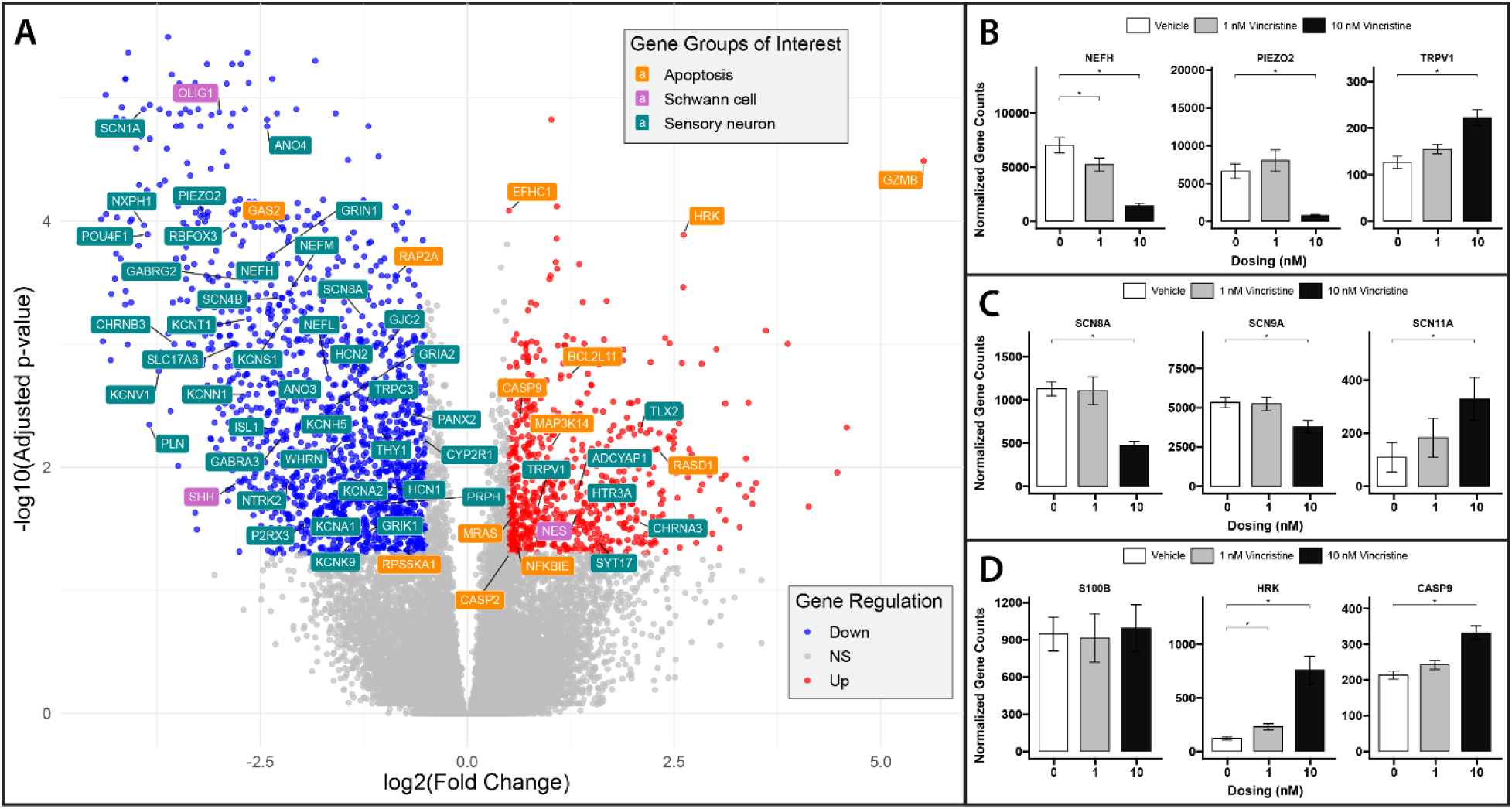
Transcriptomic profiling of vincristine exposure. (A) Volcano plot comparing bulk RNA sequencing data from vehicle- and 10 nM vincristine-treated samples (*N* = 18 wells per condition). Differential gene expression analysis was performed using DESeq2, with the x-axis representing the log₂ fold change between vincristine and vehicle groups, and the y-axis representing the –log₁₀(*p*-value). Genes with significant differential expression (*p* < 0.05) and absolute log₂ fold change ≥ 1 are highlighted: upregulated genes in red and downregulated genes in blue. Selected genes of interest are color-coded by functional category: apoptosis, Schwann cell identity, or sensory neuron markers. The data reveal broad downregulation of sensory neuron-associated genes, minimal changes in Schwann cell-related genes, and upregulation of apoptosis-related transcripts, highlighting a pattern of sensory neuron loss and stress-induced signaling following vincristine treatment. (B–D) Gene-level comparisons further support these findings. Expression of three sensory neuron markers - Neurofilament heavy chain (NEFH), Piezo2, and TRPV1 - shows significant downregulation of NEFH and Piezo2, whereas TRPV1 expression increases, consistent with stress-related hyperexcitability. Voltage-gated sodium channel genes SCN8A (Nav1.6) and SCN9A (Nav1.7) exhibit significant downregulation, further indicating functional impairment in sensory neuron excitability, while SCN11A (Nav1.9) is upregulated, again consistent with stress-associated transcriptional responses. Schwann cell marker S100 remains unchanged between conditions, suggesting a lack of major transcriptional effect on Schwann cells at this dose and timepoint. In contrast, apoptosis-related genes HRK and CASP9 are significantly upregulated, reinforcing the observation of increased cell death signaling under vincristine treatment.

## Discussion

This study introduces a more advanced Nerve-on-a-Chip model, PNS-3D organoids, as a novel human-cell-based 3D microphysiological system designed to model the PNS with endpoints analogous to clinical electrophysiological metrics, demonstrating facilitate improved human translation (Figure 1). Our findings demonstrate the system’s capacity to recapitulate key features of peripheral nerve function, including long-distance neurite outgrowth, physiological axon-glia co-development, and stimulus-evoked population level electrophysiological responses. Through integration of microelectrode arrays embedded in custom multiwell plates, we achieve scalable, high-content functional endpoints and derive clinically translatable metrics with high reproducibility (Figure 2)^1^. In addition, the system enables longitudinal, multiplexed investigations leveraging image and gene expression-based endpoints to empower deep mechanistic analysis.

Unlike conventional 2D *in vitro* models, which typically do not support densely aligned axonal bundles over extended distances or 3D myelination, our model enables both biomimetic Schwann cell alignment and, most importantly, lamellar myelin formation around axons, a hallmark of peripheral nerve physiology (Figure 3)^29^. Rodent-based 3D systems such as tissue explants and our 3D nerve-on-a-chip have shown potential as assays of peripheral myelination, but their translational value is limited by interspecies differences in axon-glia signaling and pharmacodynamics^30,31^. Our previous work with our Nerve-on-a-Chip model was the first to demonstrate human Schwann cells forming compact myelin around human peripheral axons^17^. Others have since shown evidence of myelin formation in 2D human cell-based PNS preparations^11,32^. The improvements described herein further advances the clinical translation of human-specific peripheral neurotoxicity pathways helping overcome the barrier of interspecies differences. Furthermore, unlike conventional MEA systems that use dense 2D grids to measure local field potentials or spontaneous network activity, our system features electrodes at fixed longitudinal intervals along a dense neurofiber tract capable of measuring a summation of electrophysiological responses throughout the 3D biology (Figure 4)^33^. This uniquely supports the derivation of functional population-level human-relevant readouts that mirror those used by clinicians: nerve conduction velocity and compound action potential amplitude reflecting neurological nerve conduction studies and velocity density index (VDI) representing a global functional metric. All of these advantages are contained in a unified model utilizing a multiwell plate format compatible with high-throughput workflows.

As a model for peripheral neuropathy, PNS-3D organoids enable a uniquely spatiotemporal, multiparametric assessment of neurotoxicity and neuropathy progression compared to *in vitro* or *in vivo* studies. When applied to a vincristine-induced neuropathy model, the system captured dose and time dependent deficits in nerve function with high sensitivity, yielding IC50 values that differentiated functional decline in VDI (1.23 nM) from structural changes in Degeneration Index (9.81 nM) at DIV 7 (Figures 5 and 7). These results almost directly parallel findings from clinical studies with Vincristine exposure^34,35^. In two publications across 15 studies, Cmax values are reported as free drug concentrations from 1.63 to 1.72 nM, accounting for serum levels of Vincristine of 6.52 and 6.869 nM with a 75% protein binding ratio. However, previous publications using 2D *in vitro* demonstrated a 100 fold difference in reported IC50 values. Across 100 published IC50/GC50 studies from different 2D *in vitro* based models, the average value was 173 nM. Relying too heavily on such results could have led to incorrect conclusions or discrepancies in decisions made for future work^36^. Studies in rodent models reported results within a 10-fold difference of clinical values, with significant neuropathic pain detected at 8.25 nM (average of 165 behavioral studies) in rats and 5.48 nM (average of 36 behavioral studies) in mice. Still, such discrepancies are consequential and constraints to throughput, high costs, and lack of human translation are driving the recognized need for alternative to historical animal models. Our model addresses these limitations while unlocking additional high-content endpoints and enabling multiplexed data which is critical for mechanistic understandings of biological response.

Through deeper electrophysiological analysis in a separate experiment, we can also elucidate that IC50s calculated through NCV (.73 nM) are distinct from those of CAP Amplitude (7.92 nM). In addition, spatially dependent deficits highlight additional mechanistic insights when analyzed for IC50 according to VDI binned into slow, medium, and fast conduction velocities (Figure 6). Medium and fast velocities show significant IC50 drops to 10.1 NM and 15.7 nM respectively after 1 day of VN exposure with slow velocities at 32.3 nM. After 3 days, IC50s for slow velocity decrease to 8.34 nM while medium and fast see a sharper fall to 1.55 nM and 1.5 nM. After 7 days of exposure, medium velocity VDI drops after 1 nM dosing (2.86 nM IC50) while slow and fast have already seen deficits appear before .1 nM (IC50 of .23 nM for slow and .1 nM for fast). Finally, data from spatial changes highlight that distal fibers are affected preferentially with IC50 values after 1 day of dosing of 3.02 nM distally and 22 nM proximally. However, after 7 days, both show high levels of toxicity with IC50s of .16 and .4 nM respectively. This again correlates with clinical reports which found axonopathy vs neuronopathy to be the primary mechanism of toxicity^34,35^. Taken together, results from the PNS-3D organoids reinforce the model’s ability to uniquely predict clinical results and detect early electrophysiological dysfunction before observable structural deterioration, coupled with velocity and spatially specific toxicities, offering a critical window for elucidating mechanistic insight and investigating therapeutic intervention.

Additionally, transcriptomic profiling confirmed neuron-specific degenerative pathways with preservation of Schwann cell-associated gene expression, reinforcing the model’s resolution of cell-type-specific responses (Figure 8). These molecular findings align with the observed physiological decline and suggest a progressive sensory neuron-specific degeneration with minimal Schwann cell involvement consistent with literature findings, mediated in part by activation of apoptotic pathways including HRK and CASP914^37,38^.

Together, these features represent a significant advance over rodent models and traditional 2D cultures, which lack both the human-specific context and clinically relevant readouts necessary for predictive neurotoxicity screening.

PNS-3D organoids offer significant advantages in physiological relevance and scalability, though certain limitations remain. The current system lacks vascular and immune components, which may contribute to the etiology of neurotoxicity or neuroinflammation *in vivo*. Additionally, while sensory neuron and Schwann cell co-cultures provide a robust model of the peripheral nerve, not all cell types or axonal subtypes are present, and iPSC-derived neurons may exhibit limited maturation, even after 7 weeks in culture. Electrophysiological readouts, while clinically analogous, may also be influenced by variability in organoid size and axonal density or alignment, necessitating careful standardization across experimental replicates. These limitations suggest areas for further investigation.

In summary, PNS-3D organoids occupy a critical niche among neuro-MPS tools, providing an integrated, scalable model ideally suited for safety testing and drug discovery in CIPN and beyond. It’s integration of imaging, omics, and electrophysiology supports a broad range of applications from mechanistic interrogation to high-throughput screening of therapeutic candidates.

### Future Directions

Ongoing efforts aim to further expand the platform’s translational utility. First, we are broadening compound screening for model qualification to include diverse chemotypes such as antibody-drug conjugates and targeted small molecules. Second, we are actively developing pathophysiological models for chronic pain and hereditary neuropathies, where human-relevant nerve models remain an unmet need. Lastly, enhancements to the EEA system, including microfluidic perfusion, higher-density recording, faster acquisition times, and real-time signal processing are ongoing, supporting greater throughput and more dynamic monitoring.

---

**Supplemental Figure 1.**
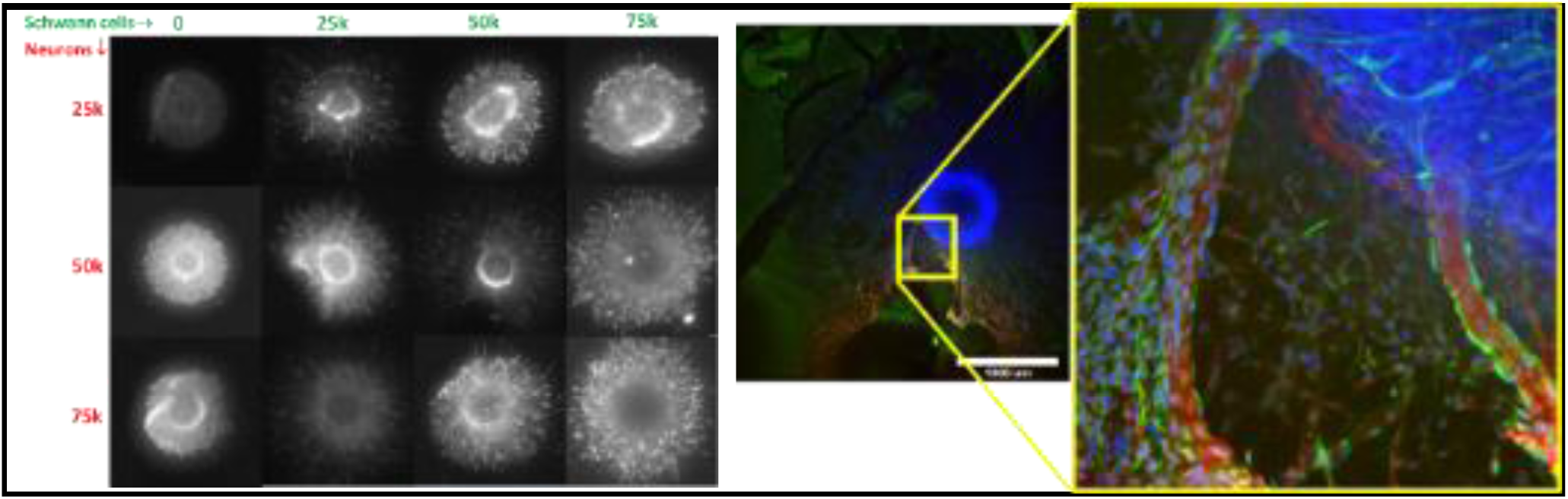
Organoid density optimization. (A) Varying Neuron and Schwann cell organoid densities were grown in 3D and stained with Beta III Tubulin, S100, and DAPI to visualize outgrowth and size. (left) A close up image (right) shows neurite outgrowth (red) Dapi (blue) and Schwann cells (green). Schwann cell alignment with neurons is critical for electrophysiological activity.

## Methods Section

### Cell culture preparation

All cell culture flasks were pre-coated with 5mL of 10µg/mL Laminin (Sigma-Aldrich, L2020) for T25 flasks (Cell Treat, 229331) and 10mL of 5µg/mL Laminin (Sigma-Aldrich, L2020) for T75 flasks (Corning, 353136). Each flask was coated under a biological safety cabinet and incubated for minimum 8 hours at 4°C temperature. After coating, the laminin solution was removed and before seeding, the plates were equilibrated with 4 mL and 9mL of culture medium to 37° in the incubator with 5% CO_2_ and a humidified atmosphere. Custom EEA plates were sterilized and coated with 0.01% Polyethyleneimine (PEI) (Sigma-Aldrich, 181978) in 1X Borate Buffer (Thermo Fisher, 28341) for 1 hour at 37°C temperature. After the incubation period the plates dried for 8 hours and were subsequently coated with 10µg/mL Laminin solution in PBS (Caisson Labs, PBL06) and incubated for minimum 8 hours at 4°C temperature. After coating, the laminin solution was removed. Before seeding the coculture organoids, the plates are equilibrated with 200µL per well of 10µg/mL Laminin supplemented Pre-myelination culture media in the 37° in the incubator. Pre-myelination cell culture media consisted of Neurobasal Medium (Gibco, 21103049) supplemented with 2% B27, (Gibco, 175040044) 1% Glutamax, (Gibco, 35050061) 1% Antibiotic-Antimycotic, (Sigma-Aldrich, A5955) 20ng/mL Human Nerve Growth Factor, (Peprotech, 450-01) 10ng/mL Human Brain Derived Neurotrophic Factor, (Gemini Bio, 300-107P) 10ng/mL glial cell line derived neurotrophic factor, (Peprotech, 450-10) and 10% filtered fetal bovine serum. (Hyclone, SH30022.0)

### Organoid Fabrication

Ampules of primary human Schwann cells (hSCs) were briefly thawed at 37°C. The contents of the vials were transferred into a conical tube containing 9mL of chilled Schwann cell Low Proliferation media. (LPM) LPM consist of filtered Dulbecco’s modified eagle medium, (Hyclone, SH30022.0) supplemented with 10% decomplemented fetal bovine serum (Hyclone, SH30396.03), 1% Glutamax, (Gibco, 35050061) and 50mg/mL gentamicin (Sigma-Aldrich, G1397). FBS was decomplemented in house by warming aliquoted FBS at 50°C for 1 hour. The cell suspension was immediately centrifuged at 200 RCF for 10 minutes at 4°C to form a cell pellet. The media was aspirated, and the pellet was resuspended and gently triturated with a pipette. 10µL was collected for counting and viability. Cells were counted with the Nexcelom Cellometer K2 and stained with Nexcelom Viastain AO/PI solution (Nexcelom, CS2-0101). The cell pellet was then resuspended in 10mL of Schwann cell high proliferation culture media (HPM) and plated in the precoated T75 flask. HPM consists of LPM supplemented with 2µM forskolin (Sigma-Aldrich, F6886) and 10nM human Heregulin β1. (Peprotech, 100-03) The Schwann cells were incubated at 8.5% CO2 in a humidified environment for up to 48 hours before passaging to allow for proliferation. To passage, hSCs were rinsed with Hank’s balanced salt solution (HBSS) (Sigma, 55021C) and passaged with 0.05% trypsin, 0.02% EDTA (Sigma-Aldrich, 59408C) for five minutes at room temperature. The hSCs were collected and centrifuged for 10 minutes at 200RCF and 4 °C, the pellet was rinsed, resuspended and centrifuged again at the same speed and time. When the cells are pelleted, the excess media is removed and the cells are counted for organoid generation. Ampules of RealDRG™ hIPSC derived sensory neurons (hSNs) sourced from Anatomic Incorporated (1020-1M) were briefly thawed at 37°C. The contents of the vials were transferred into an empty conical tube. The vial was rinsed with 1mL of Anatomic SensoMM culture media (Anatomic Inc, SensoMM) and additional SensoMM is dispensed into the conical for a total of 5mL per vial thawed. The cell suspension is immediately centrifuged at 300 RCF for 4 minutes to form a cell pellet. The media was aspirated, and the pellet was resuspended with 1mL of culture media and gently triturated with a pipette. 10µL was collected for counting and viability. Cells were counted with the Nexcelom Cellometer K2 and stained with Nexcelom Viastain AO/PI solution. The cell suspension was transferred to a precoated and pre-equilibriated T25 flask. The cells incubatedin a humidified environment at 5% CO2 and 37°C for no more than 18 hours before passaging. To passage, culture media was removed hSNs were rinsed with PBS and lifted from the flask with Accumax (Sigma-Aldrich, SCR006) incubated at 37° for 7 minutes. The cells were collected and centrifuged for 4 minutes at 300RCF, the pellet was rinsed and centrifuged again at the same speed and time. Both hSCs and hSNs were passaged on the same day prior to forming coculture organoids. Each cell pellet was resuspended in 1mL of SensoMM culture media and 10µL was collected for counting and viability. Cells were counted with the Nexcelom Cellometer K2 and stained with Nexcelom Viastain AO/PI solution. hSCs and hSNs were diluted to the appropriate cell density requirement to create a cell suspension of 50,000 hSNs and 25,000 hSCs per 200µL of SensoMM. 200µL of cell suspension was plated in a round bottom 96-well plate (Thermo Fisher, 74929) and centrifuged for 4 minutes at 300RCF to create the organoid. The organoids bundled and compacted over 4 days with a half media change at day 2 with SensoMM. On day 4, organoids were imaged for size analysis metrics and placed in PNS-3D organoid cultures.

### PNS-3D organoid fabrication and growth

The organoids were placed into the top bulb of the EEA plate using micro-forceps (Dumostar, 11294) and a microscope in a Thermo Fisher-Scientific Heraguard Eco Clean bench. Individual organoids were initially transferred from the 96-well U-bottom plate into a sterile 100mm petri dish under the BSL2 safety cabinet. The petri dish and lid was transferred to the clean bench where the equilibrated EEA plate was stored. The warmed laminin-supplemented media was removed, and the organoids were transferred with micro-forceps from the petri dish to the top bulb of the EEA channel. The organoids were covered in laminin supplemented Pre-Myelination media and returned the incubator for a minimum of one hour to ensure adherence to the plate before the laminin supplemented media was removed and replaced with Pre-myelination media. The organoids adhered for 3 days undisturbed in the incubator; on day 3 the entirety of the channel was covered in Matrigel (Corning, 354230) to allow for 3D growth over the electrodes. The matrigel was placed by removing the cultures from the incubator and placing it in the clean bench. Culture media was removed with a pipette to expose the channel. 2-3µL of Matrigel was dispensed into bulb without the organoid, filling and covering the entirety of the channel. After 2-3 minutes, once the matrigel was hardened 200µL of fresh Pre-myelination media was dispensed into each well. After this matrigel procedure and media change, a regular Monday, Wednesday, Friday full 200µL media change was established for the following 7 weeks. Several media types were used; for the first week, Pre-myelination media was administered to the culture, for weeks 2-5 Myelination media (Neurobasal Medium supplemented with 2% B27, 1% Glutamax, 1% Antibiotic-Antimycotic, 20ng/mL Human Nerve Growth Factor, 10ng/mL Human Brain Derived Neurotrophic Factor, 10ng/mL glial cell line derived neurotrophic factor, 10% filtered fetal bovine serum, and 50µg/mL Ascorbic Acid (StemCell, 72132)) was administered and for week 6 Brainphys Myelination Media (Brainphys Media, (Stemcell, 5790) supplemented with 2% B27, 1% Glutamax, 1% Antibiotic-Antimycotic, 20ng/mL Human Nerve Growth Factor, 10ng/mL Human Brain Derived Neurotrophic Factor, 10ng/mL glial cell line derived neurotrophic factor, 10% filtered fetal bovine serum, and 50µg/mL Ascorbic Acid.)) Weekly imaging was performed during the growth weeks to ensure PNS-3D organoid cultures were healthy and robust before dosing took place.

### Dosing

While undergoing compound testing in the final week, a variation of media containing no serum was administered to the culture to prevent any neuroprotective effects of serum. Brainphys dosing media consisted of Brainphys Media supplemented with 2% B27, 1% Glutamax, 1% Antibiotic-Antimycotic, 20ng/mL Human Nerve Growth Factor, 10ng/mL Human Brain Derived Neurotrophic Factor, 10ng/mL glial cell line derived neurotrophic factor, 10% filtered fetal bovine serum, and 50µg/mL Ascorbic Acid. Cultures were dosed on a Monday, Wednesday, Friday for a total of seven days of exposure to Vincristine. 1X dilution of each concentration of Vincristine (1nM, 10nM, and 100nM) was made on Monday, Wednesday, and Friday in Brainphys Dosing Media before the media change was performed.

### Electrophysiological Data Collection

During the week before dosing and dose week several rounds of stimulated electrophysiology were performed in a designated incubator with 5% CO_2_. 5 days of limited electrode stimulation was done prior to the addition of test compounds. Before applying the compounds, a baseline reading of full electrode stimulation was recorded. Dosing proceeded, mid-week a limited electrode stimulation was recorded, and on the last day of culture and 7 days of compound testing the last baseline recording was measured. Electrophysiology was performed before media changes to ensure a fresh media change would not affect the outcome.

A custom MATLAB R2024a application was written to communicate with Intan RHX v3.1.0 in real time via TCP/IP, automating stimulation and acquisition for a single 24-well plate. Stimulation parameters were read line-by-line from an Excel “stimulation script,” which specified the current amplitude, stimulation electrode, pulse width, polarity, trial frequency, number of trials, and number of pulses per trial for each stimulus set. During a typical session the pulse width was fixed at 0.1 ms, the polarity at cathodic-first biphasic shape, trial frequency at 1 Hz, number of trials at 12, and number of pulses per trial at 1. Each session delivered twenty-five distinct stimulus sets: five current levels (1, 5, 10, 24, 48 µA) applied to five stimulation sites (three proximal [<2.5 mm from organoid], two distal [>2.5 mm from organoid]), yielding approximately 250 recordings per well (10 recording electrodes × 25 sets).

Electrically evoked recordings were delivered through current-controlled, cathodic-first biphasic pulses with matched phase amplitudes and durations to ensure charge balance. For each stimulation set, the MATLAB controller issued TCP commands to the Intan RHX software, programming the on-board RHS2116 stimulators with the specified current, polarity, and pulse width, then triggering acquisition and precisely timing stimulus delivery to the designated electrodes.

Hardware filters on the amplifier were configured with an analog passband of 80–7500 Hz and a digital offset removal cutoff of 100 Hz to suppress mains interference while preserving compound action potential (CAP) content. During each stimulus the Intan amplifiers temporarily switched to a reduced bandwidth (lower cutoff ≈ 1000 Hz) to minimize stimulus artefact magnitude. The sampling rate for the recording was 30kHz.

Raw data were saved by the RHX software in binary (*.rhd*) format and processed offline in MATLAB. Continuous files (∼12 s per configuration) were segmented into individual trials using the timestamp of each stimulus onset, producing a time-aligned matrix (time on x, trials on y) cropped to −40 ms to +60 ms relative to the stimulus. Low-frequency stimulus drift was removed by subtracting a 10 Hz low-pass-filtered version of the matrix (filtfilt + Butterworth, 4th order) from the original, effectively eliminating artefact tails without distorting CAP features within ∼1.5–2 ms of stimulus offset. Baseline-corrected trials were then averaged to enhance signal-to-noise ratio prior to metric extraction.

### Imaging methods

Organoid images were taken on the Molecular Devices ImageXpress Micro Confocal Microscope on DIV 4 of the organoid formation phase before organoids were placed in the bulb of the EEA channel to ensure all organoids had formed correctly. The TL25 images were taken with a 4X objective with a 0.05ms exposure time. Weekly imaging was performed once per week to ensure the health and growth of the cultures. Cultures were imaged with the Molecular Devices ImageXpress Micro Confocal Microscope in TL25, with a 10X objective for 7.27ms exposure. Z planes of 19 steps across 8 sites down the channel allowed us to visualize the 3D nature of the peripheral nerve. The range was 15µm between steps for a total range of 270µm. During the dosing week, cell cultures were imaged 2 additional times, once in the middle of the week and once at the end of the seven day dose period. Images were stitched together with a custom Matlab application for growth analysis and monitoring.

### Staining methods

Immunohistochemistry was performed on DIV49 cultures; Beta III Tubulin (neurite outgrowth) and S100 (Schwann cells) allowed us to visualize the extent of cell migration and Schwann cell alignment with neurites at the completion of the culture period. Cultures were washed with PBS half changes 3 times and the plates were subsequently fixed for 30 minutes in a 4% PFA (EMS, 15710) solution. After the incubation period the PFA was washed with PBS half changes 4 times. Cultures were blocked for one hour in a solution consisting of PBS, 4mg/mL BSA, (Sigma-Aldrich, A2153) 0.2% TritonX, (Sigma-Aldrich, X100) and 5% Normal Goat Serum (Jackson ImmunoResearch, 005-000-121). The blocking solution was furthered used for secondary antibody incubation. The primaries included were a ready to use rabbit-anti-S100 (DAKO, GA50462-1) and 1:100 dilution of mouse-anti-Beta III Tubulin (Abcam, ab78078) were incubated overnight, the plates were washed 6 times with PBS half changes. The secondary stains were each a 1:250 dilution, goat-anti-rabbit 647 (Thermo Fisher, A32733TR) and goat-anti-mouse 488 (Abcam, 150113) were applied for a 75-minute incubation at room temperature before being washed out 6 times with PBS half changes.

### Fluorescent microscopy

Fluorescent Z-stack images were acquired with a Nikon SMZ25 top-down stereo microscope fitted with a motorized XYZ stage and motorized filter wheel. A top-illumination geometry was required because the polyimide film at the base of the EEA plates blocks inverted-path fluorescence. Images were captured at 20X optical zoom with a 1 s exposure for each channel. For every field of view, a Z-series of 30 optical sections was collected at 25 µm intervals (total depth ≈ 0.75 mm), ensuring the entire PNS3D organoid volume was sampled. Acquisition parameters—including stage coordinates, filter selection, exposure time, and Z-step spacing—were scripted in NIS-Elements for unattended, plate-wide imaging. Raw images were saved as 16-bit TIFF stacks.

### Snap freezing for transcriptomics

The snap-freezing bath was prepared by filling a large rectangular ice bucket with enough methanol (Sigma-Aldrich, 179337) to create a layer approximately ½ inch deep, sufficient to cover an EEA plate. Dry ice pellets were added piece by piece until temperature equilibrium was reached, indicated by the pellets no longer melting rapidly. Plates were prepared and snap frozen one at a time. For each plate, 200µL of media was removed from each well, followed by three washes with 200µL of PBS. The final PBS wash was carefully removed to ensure no residual PBS remained, as this was critical for effective snap freezing. A plate seal was placed over the wells, the lid was returned to the plate, and the plate was securely wrapped with parafilm. The sealed plate was then placed into a properly sized plastick zip bag. Using tongs, the bagged plate was slowly submerged into the dry ice-methanol bath. The plate was allowed to sit in the bath for at least one minute, during which the tongs were used to gently push the plate down to prevent floating; Matrigel turned white upon freezing. The plate was then removed from the bath and checked to ensure there were no cracks and that no methanol had entered the wells. The snap-frozen plate was placed on dry ice and transported to the -80 °C freezer for storage. If additional plates were processed, the dry ice in the bath was replenished as needed. Total RNA was extracted using the Quick-RNA™ Miniprep Kit (Zymo Research, R1054) according to the manufacturer’s instructions.

### TEM methods

To capture transmission electron microscopy of sliced cross-sections of the peripheral nerve model, a previously developed modality was utilized in which a key-hole construct was fabricated, enabling preservation and embedding of a fully formed peripheral nerve model. A dual-hydrogel scaffold was prepared on the membranes of Transwell® inserts (0.4 µm/PES; Corning, 3450) using a micro-photolithography technique. All solutions were created with sterile-filtered PBS unless otherwise noted. The outer cell-restrictive (i.e., growth-resistant) photo-translinkable hydrogel was created using a solution of polyethylene glycol dimethacrylate 1000 (PEGDMA; Polysciences, 15178) and lithium phenyl-2,4,6-trimethylbenzoylphosphinate (LAP; Tocris, 6146). First, 30% w/v PEGDMA and 1.1 mM LAP solutions were created and mixed in a 1:1 solution. The resulting 15% solution was sterile-filtered and stored for 8 hours at 4°C. The following day the 600µL of the solution was added to Transwell® inserts while positioned under the lens of a Digital Micromirror Device (DMD, PRO4500 Wintech Production Ready Optical Engine; Wintech Digital Systems Technology Corp, Carlsbad, CA) on Rain-X (ITW Global Brands, Glenview, IL)-treated glass slides. The mask and polymerization parameters were selected using commercial software (DLP Lightcrafter 4500 Control Software, Texas Instruments, Dallas, TX), and irradiation of the photo-translinkable solution occurred for 60 seconds using the ultraviolet light of 385 nm wavelength. After treatment, excess PEGDMA/LAP solution was removed from the insert and from within the void created by the photomask. The constructs were then washed using 2% Antibiotic/Antimycotic wash buffer in PBS three times on the top and bottom of the insert for 10 minutes each. Wash buffer was removed from the insert and the inner keyhole-shaped channels were exposed before placing DIV 4 organoids into the keyhole.

### Transwell culture methods

DIV4 organoids were placed into each Transwell construct as previously described in PNS-3D organoid fabrication and growth. The organoids were then covered in Matrigel and 1.5mL of Pre-myelination media was placed below the Transwell to diffuse across the membrane and into the culture on Monday, Wednesday, and Friday for seven days. For the next three weeks media changes happened three times per week with Myelination media. On DIV 28 cultures were fully matured and growth reached the end of the channel. Cultures underwent weekly imaging as described above to monitor growth and health.

### Plastic resin embedding

All materials used for embedding were purchased from Electron Microscopy Sciences unless otherwise noted and were handled under a chemical flow hood with recommended personal protective equipment. Plastic embedding is essential for evaluating the ultrastructure of the axons which is difficult to quantify by just performing immunostaining. AT DIV28, the hydrogel constructs were washed 3 times with PBS before fixation. To fix, the hydrogel constructs were soaked in a solution of 4% PFA/2% glutaraldehyde (EMS, 16221) in 0.1M sodium cacodylate (EMS, 11650) with 3% sucrose (BioBasic, SB0498) and 3mM CaCl_2_ (Sigma-aldrich, 449709) for 3 hours at room temperature. After fixation constructs were washed 4 times with PBS.

With a scalpel, hydrogel constructs were dissected individually from the transmembrane wells, without removal of the PEGDMA and embedded on a glass slide in a 3% Agarose (Sigma-Aldrich A9539) in distilled water solution and allowed to harden over ice. Once the gels were solid they were transferred to glass jars for the remainder of the process. Secondary fixation and staining of cellular lipids was achieved by post-fixation with 2% osmium tetroxide (EMS, 19150) in PBS, pH 7.4, for 2 hours under dark conditions at room temperature. The constructs were then washed with water 3 times. Counterstaining was performed with 1% aqueous uranyl acetate (EMS, 22400-2) overnight on a rocker. The next day uranyl acetate solution was washed three times with distilled water for ten minutes each.

Dehydration was done with graded ethanol (EMS, 15056) washes at room temperature. Dehydration started with 30% ethanol, followed by incubation on a rocker for 20 minutes at room temperature. The 30% ethanol was then replaced with 50% ethanol, and the vials were incubated again for 20 minutes at room temperature. Finally, the 50% ethanol was removed and replaced with 70% ethanol, and the vials were incubated on the rocker overnight at room temperature.

The dehydration process continued the next day by removing the 70% ethanol and adding of 80% ethanol per vial. The vials were then incubated on a rocker for 15 minutes. After incubation, the 80% ethanol was removed and replaced with 90% ethanol per vial, and the vials were again incubated on the rocker for 15 minutes. This step was followed by replacing the 90% ethanol with 100% ethanol per vial, with a 15-minute incubation on the rocker. The 100% ethanol incubation was repeated two additional times for a total of three 15-minute incubations.

Finally, the 100% ethanol was removed and replaced with 10 mL of propylene oxide (EMS, 20401) per vial. The vials were incubated on the rocker for 15 minutes, and this propylene oxide incubation was repeated once more for a total of two 15-minute incubations.

The propylene oxide was removed from each vial and replaced with 2 mL of a 1:1 resin to propylene oxide (PO) solution. The vials were incubated on a rocker for 2 hours at room temperature. After incubation, the 1:1 resin:PO solution was removed and replaced with 2 mL of a 2:1 resin:PO solution, followed by another 2-hour incubation on the rocker at room temperature. Finally, the 2:1 resin:PO solution was removed and replaced with 2 mL of 100% resin, and the vials were incubated overnight on the rocker at room temperature. The following day, the constructs were moved to new vials containing fresh 100% resin and incubated for four hours. The constructs were then transferred to Flat Embedding Molds (EMS 70902) and filled with 100% resin and cured for 48 hours at 65°C.

### Sectioning and transmission electron microscopy (TEM)

Sectioning and TEM evaluation was performed at the Shared Instrumentation Facility (SIF) at Louisiana State University (Baton Rouge, LA). Ultrathin sections were cut to a thickness of 80–100 nm at four locations within the PNS-3D organoid specimen: within the bulb of the tissue, where the bulb met the channel and the proximal channel (i.e., near the bulb), and distal channel. Sections were placed on Formvar carbon-coated copper grids, 200 mesh, and impregnated with metal by floating on droplets of 2% uranyl acetate for 20 mins at RT. They were then rinsed with deionized water droplets 3 times, for 1 min. To visualize, a JEOL 1400 TEM (Peabody, MA) was used with an accelerating voltage of 120 kV at varying magnifications.

### Data analysis

#### Organoid size analysis

Bright-field images of DIV4 organoids cultured in round-bottom 96-well plates were analyzed to ensure consistency of each batch with a custom MATLAB R2024a application that automated image import, processing, and metric export. Each image underwent median filtering and background subtraction, followed by adaptive thresholding to isolate the organoid from the background. The resulting edge trace was convered into a binary mask that was refined with imfill and bwareaopen to close internal holes and eliminate small debris. Features were extracted with regionprops, and two size-related metrics were reported for every organoid: the *equivalent diameter*, defined as the diameter of a circle with the same area as the binary mask, and the *solidity*, calculated as the ratio of the mask area to its convex-hull area, providing a measure of compactness. All measurements were exported as comma-separated values for downstream statistical analysis in R 4.3.1.

#### Cell migration analysis

Growth was tracked using a custom MATLAB application both automatically and manually. Manually, each stitched image containing all Z-stacks were overlayed with a grid marking segments of 100µm increments. The grid begins at the bulb on the left where the organoid is placed and extended down the entirety of the channel. Because the axons (brightfield) or stained cells (IHC; Schwann cells, sensory neurons, or cell nuclei) were visible in multiple Z-stacks it was important to consider each focus point to find the longest and densest growth. Longest neurite position was determined to be the increment of grid that contained at least 3 axons or cells.

#### Electrophysiological noise analysis

A total of sixteen 24-well EEA plates were tracked longitudinally between DIV35 and DIV86. On each recording day, plates were transferred to the isolated electrophysiology incubator and connected to the custom headstage and Intan RHS system to measure electrophysiology under standard tissue culture settings (37 °C, 5% CO2). Evoked stimulation trials (n = 12 per well and timepoint) were delivered as described in the Electrophysiological Data Collection section. Immediately prior to each stimulus, a 20 ms baseline segment was isolated from the raw voltage trace using custom MATLAB scripts. Two noise metrics were calculated for every electrode across all wells and plates. Root-mean-square noise (Vrms) was determined as 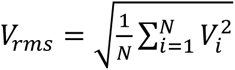, where N is the number of samples in the 20 ms window. Maximum peak noise (Vpeak) was defined as the largest absolute voltage value observed within the same window, *V*_*peak*_ = max {|*V*_*i*_|}. Vrms and Vpeak values were exported as .csv files and aggregated across trials in R 4.3.1 using dplyr 1.1.4 before plotting using the ggplot2 package.

#### Maximum Velocity Projection and Velocity Density Index Calculation

The ten recording electrodes and five stimulation sites used in each recording yielded forty-five unique recordings (12 trials each at 1 Hz) for each PNS-3D organoid (ten electrodes × five sites minus the five recordings at the stimulation site). Trial-averaged voltage traces were converted from the time domain to a conduction-velocity domain by dividing the recording-electrode/stimulation-electrode separation distance (*d*, mm) by the time vector (*t*, ms). The analytic signal envelope was then obtained with MATLAB’s hilbert function to produce a positive-valued waveform of voltage versus velocity. All forty-five envelopes were resampled with cubic interpolation (interp1, velocity increment = 0.01 m s⁻¹) to allow alignment in velocity space. At each velocity breakpoint, the maximum value across traces was calculated to form the maximum-velocity-projection (MVP) waveform. The Velocity Density Index (VDI) was calculated as the trapezoidal area under the MVP between 0 and 1.5 m s⁻¹ above a threshold of 5 uV to eliminate noise.

Variants of the analysis were generated by selecting only proximal (< 2.5 mm) or distal (> 2.5 mm) stimulation conditions (spatial binning) or by integrating the MVP over predefined velocity bands—slow (0–0.5 m s⁻¹), medium (0.5–1 m s⁻¹), and fast (1– 1.5 m s⁻¹)—to assess how different conduction regimes evolved with time and pharmacological treatment.

All calculations were performed in MATLAB R2024a. Data collation, organization, and visualization were carried out in R 4.3.1 using dplyr 1.1.4 and ggplot2 3.5.1.

#### CAP detection and metric calculation

A three-layer bidirectional long short-term memory (biLSTM) neural network (250, 100, and 50 hidden units) with 0.5 dropout between layers and a final softmax classification layer was implemented in MATLAB R2024a (Deep Learning Toolbox) to label 0–20 ms post-stimulus segments as *compound action potential (CAP)*, *noise*, stimulus artefact, or *no response*. The network was trained on 765 curated recordings (265 CAP, 250 noise, 250 blank) using a 70 %/30 % train/validation split and proportional class weights to mitigate class imbalance. Input tensors contained the full trial set, the across-trial average, and five sliding-window statistics (window = 5 samples): RMS mean, standard deviation, maximum, minimum, and range. The trained model achieved 91.87 % accuracy on the hold-out validation set.

Batch inference of the classifier produced time stamps for every detected CAP segment. For each recording electrode–stimulus electrode–current–timepoint combination the following metrics were derived:

- **Nerve-conduction velocity (NCV, m s⁻¹)** : distance between electrodes (mm) divided by latency from stimulus onset to peak absolute voltage (ms).
- **Amplitude (µV)** : peak-to-peak voltage (maximum − minimum) of the CAP segment.
- **Response count (#)** : number of CAP segments detected within 0–20 ms.

When multiple CAPs were present in a recording, NCV and amplitude values were averaged across segments. Metrics were then aggregated across replicates to yield sample-level means for each time-point and current, which served as inputs for normalization and dose– response modelling.

#### Electrophysiology data normalization and dose-response modeling

Baseline values for each electrophysiological metric – nerve conduction velocity (NCV), compound action potential (CAP) amplitude, Velocity Density Index (VDI), number of responses, velocity-binned VDI, and spatially binned VDI - were recorded immediately prior to compound exposure. Post-dosing measurements were acquired at one or more subsequent time-points (e.g., 1 day, 3 days, 7 days, etc). For every sample-metric pair, non-baseline readings were first normalized to the sample-specific baseline (metric/metric₀ × 100 %). These values were then corrected for vehicle effects by dividing by the mean vehicle response at the corresponding time-point and scaling again to 100 %, enabling direct comparison of drug-induced changes.

Dose–response (IC₅₀) curves were generated from the mean response of at least six biological replicates per dosing condition. Curves were fitted with the drc package (v3.0-1) in R 4.3.1 using a five-parameter log-logistic model (LL.5) with the upper asymptote constrained to 100 % and the lower to 0 %. The remaining three parameters (slope, inflection point, and asymmetry) were estimated by nonlinear least squares, and IC₅₀ values were extracted from the model output.

Statistical significance between vehicle and treated groups at each time-point was assessed with two-sided *t*-tests (stats::t.test) assuming unequal variances where *p* < 0.05 was considered significant. All plots were produced with ggplot2 3.5.1, displaying the mean ± standard error of the mean (SEM), individual data points where appropriate, fitted IC₅₀ curves, and significance annotations.

All data normalization and visualization workflows were scripted in R with dplyr 1.1.4, tidyr 1.3.0, and ggpubr 0.6.0.

#### Brightfield image processing

Bright-field Z-stack images were imported into MATLAB R2024a, and eight regions of interest (∼900 µm x ∼150 µm) were cropped along the channel in 1 mm distance between electrodes to avoid sharp edges and retain only cellular features. Each region contained 18 optical sections (Z-steps).

Per Z-step, a custom Niblack thresholding routine was applied to the grayscale image to generate an initial binary edge mask. The mask was refined with bwpropfilt to remove objects ≤ 10 px². Properties of the remaining regions were extracted with regionprops to produce two types of binary masks. First, the fiber mask was obtained by filtering for objects with an aspect ratio ≥ 0.9 (major/minor axis) and a major-axis length ≥ 25 pixels, yielding a binary representation of axon-like fibers. Second, the fragment mask was generated by filtering for objects between 10 px² and 25 px² in area, capturing debris-like fragments.

Three morphology metrics were computed from these two masks:

1. **Fiber count** — number of regions in the fiber mask.
2. **Fiber length** — mean major-axis length of fiber-mask regions.
3. **Degeneration index** — number of regions in the fragment mask.

Metrics were calculated for every region and Z-step, producing 144 measurements per sample (8 regions × 18 Z-steps) at each time-point for downstream statistical analysis.

#### Nerve morphology data normalization and dose-response analysis

Each bright-field acquisition yielded 144 raw measurements per morphology metric (8 segments × 18 Z-steps) for every sample and time-point. To follow local changes over time, every segment/Z-step value at post-dosing time-points was first normalized to its own pre-dose baseline: *Metric*_*rel*_ = (*Metric*⁄*Metric*_*baseline*_) × 100%. The resulting relative values were then averaged across all segments and Z-steps to obtain a single sample-level metric at each time-point. Because degeneration metrics seldom reach zero, a second normalization was applied to anchor the vehicle at 100 % and a toxic reference (1000 nM vincristine at 7 days) at 0 % for each metric/time-point:

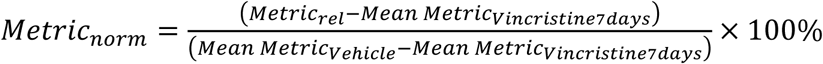

Negative values were set to 0 %. The procedure produced three interpretable metrics - fiber count, fiber length, and degeneration index - expressed as percentage deviation from vehicle.

Dose–response (IC₅₀) curves, statistical tests, and visualization were accomplished similar to the electrophysiological metrics above.

#### Differential gene expression analysis

Bulk RNA-seq libraries were generated from peripheral-nerve organoid cultures after 7 days of vincristine dosing. Twenty-seven libraries were analyzed, corresponding to three biological replicates per run across three runs for each treatment group: vehicle, 1 nM vincristine, and 10 nM vincristine (9 samples per group). Each library was prepared from RNA pooled from two PNS-3D organoids to achieve sufficient input.

Sequencing was performed by Novogene, delivering raw gene counts, which were imported into R 4.3.1. Normalization and variance-stabilizing transformation were performed with DESeq2 v1.46.0, yielding counts suitable for downstream analysis without conversion to TPM or FPKM. Log transformed expression values were analyzed with the limma v3.62.2 workflow (*lmFit* followed by *eBayes*). Differential expression between 10 nM vincristine and vehicle was summarized with *topTable*, and genes satisfying |log₂fold change| > 2 with Benjamini–Hochberg adjusted *p* < 0.05 were designated significantly up or downregulated. Pairwise comparisons (vehicle vs 1 nM, vehicle vs 10 nM) for individual genes employed two-sided *t*tests consistent with earlier statistical procedures. Resulting gene lists were visualized with ggplot2 3.5.1.

## Acknowledgements

The authors thank Lise Harbom for developing the transmission electron microscopy (TEM) protocols in this study and Ying Xiao (Louisiana State University, Baton Rouge) for carrying out the TEM imaging and further refining those methods. We thank Wesley Anderson for developing imaging methodologies and imaging of immunohistochemistry samples. We thank Diego Gatica, Danielle Roof, Madelyn Korn, and Cameron Myers for their assistance with tissue culture and data collection. We also thank Anatomic Inc. for their support in testing, characterizing, and integrating their iPSC-derived human sensory neurons into our model.

